# Wnt3 distribution in the zebrafish brain is determined by expression, diffusion and multiple molecular interactions

**DOI:** 10.1101/2020.05.29.124560

**Authors:** Sapthaswaran Veerapathiran, Cathleen Teh, Shiwen Zhu, Indira Kartigayen, Vladimir Korzh, Paul T. Matsudaira, Thorsten Wohland

**Author notes:** Author for correspondence Thorsten Wohland.

## Abstract

Wnt3 proteins are lipidated and glycosylated, secreted signaling molecules that play an important role in zebrafish neural patterning and brain development. However, the transport mechanism of lipid-modified Wnts through the hydrophilic extracellular environment for long-range action remains unresolved. Here, we determine how Wnt3 accomplishes long-range distribution in the zebrafish brain. First, we characterize the Wnt3-producing source and Wnt3-receiving target regions. Subsequently, we analyze Wnt3 mobility at different length scales by fluorescence correlation spectroscopy and fluorescence recovery after photo-bleaching. We demonstrate that Wnt3 spreads extracellularly and interacts with heparan sulfate proteoglycans (HSPG). We then determine the binding affinity of Wnt3 to its receptor, Frizzled1 (Fzd1), using fluorescence cross-correlation spectroscopy, and show that the co-receptor, low-density lipoprotein receptor-related protein 5 (Lrp5), is required for Wnt3-Fzd1 interaction. Our results are consistent with the extracellular distribution of Wnt3 by a diffusive mechanism that is modified by tissue morphology, interactions with HSPG and Lrp5-mediated receptor binding, to regulate zebrafish brain development.

## Introduction

Wnt proteins represent a family of secreted signaling glycoproteins having multiple functions in embryonic development such as specification of the vertebrate axis, embryonic induction, maintenance of cell potency, cell fate determination, cell migration, cell division, and apoptosis, to name a few (Clevers & Nusse, 2012; Hikasa & Sokol, 2013; Logan & Nusse, 2004; Moon et al., 2002). So far, 13 *wnt* gene subfamilies have been identified, although the number of *wnt* genes differs between species (Schubert & Holland, 2013). Wnts are generally 350 - 400 amino acids in length (molecular weight of ∼40 kDa), with highly conserved cysteine residues. Wnts are hydrophobic and water-insoluble due to their post-translational lipidation in the endoplasmic reticulum (ER) (Mikels & Nusse, 2006). Porcupine (Porc), an O-acyltransferase localized on the membrane of the ER, catalyzes the acylation of Wnts and provides Wnts hydrophobic characteristics (Herr & Basler, 2012). The acylation facilitates the interaction of Wnts with Wntless, a transmembrane protein that shuttles Wnts to the plasma membrane (Galli et al., 2007). From the plasma membrane, they are secreted and transported to Wnt-receiving cells. Hence, the acylation of Wnts is a critical step for their trafficking, secretion and activity (Coudreuse & Korswagen, 2007).

The addition of lipid moieties makes the long-range free diffusion of Wnts in the aqueous extra-cellular matrix problematic. Several transport mechanisms were proposed to explain how Wnts navigate the aqueous environment to achieve long-range action (Routledge & Scholpp, 2019). Facilitated shuttling of Wnts by chaperone proteins is a commonly reported mode of distribution. Here, Wnt-binding proteins such as secreted Frizzled-related proteins (sFRPs) (Esteve et al., 2011; Mii & Taira, 2009), Secreted Wg-interacting Molecule (Swim) (Mulligan et al., 2012) or afamin (Mihara et al., 2016) shield the hydrophobic regions of Wnts and provide stability in the aqueous environment. Similarly, hydrophobic Wnt molecules could be packaged inside exosomes and lipoprotein particles, which enables their extracellular movement (Greco et al., 2001; Neumann et al., 2009; Panáková et al., 2005). Additionally, heparan sulfate proteoglycans (HSPG) present in the extracellular matrix serve as binding sites for several signaling molecules, including Wnts (Kirkpatrick & Selleck, 2007). HSPG maintains the solubility of Wnt ligands, and prevents their aggregation in the aqueous extracellular matrix, thereby enhancing their range and function (Fuerer et al., 2010; Mii et al., 2017). Further evidence suggests that HSPG in coordination with Wnts are pivotal in regulating gastrulation, neurulation and axis formation during embryonic development (Ohkawara et al., 2003; Saied-Santiago et al., 2017; Tao et al., 2005; Topczewski et al., 2001). On the other hand, it was also recently noticed that certain Wnts could be deacylated by Notum, a secreted deacylase, but retain their signaling activity (Speer et al., 2019). Besides the extracellular transport mechanism, certain Wnt proteins may also reach their target tissues through filopodial extensions called cytonemes, as seen for Wnt2b in *Xenopus* (Holzer et al., 2012), Wg in *Drosophila* (Huang & Kornberg, 2015) and Wnt8a in zebrafish embryos (Mattes et al., 2018; Stanganello et al., 2015).

Finally, when Wnts reach their target tissues, they bind to their target receptors and elicit a signaling cascade. To date, Wnts are known to interact with more than 15 receptor and co-receptor protein families (Niehrs, 2012), of which the Frizzled (Fzd) receptor super-family is the most commonly investigated. Fzd proteins are categorized under the Class-F super-family of G-protein coupled receptors. The super-family comprises 10 Fzd receptors (Fzd1-Fzd10) and Smoothened (SMO) (Schulte & Wright, 2018), all with a seven-pass transmembrane domain and a highly conserved cysteine-rich domain (CRD) (Hsieh et al., 1999; Wu & Nusse, 2002). Structural studies revealed that the low-density lipoprotein receptor-related protein (Lrp-5/6) acts as a co-receptor and is involved with the Wnt-Fzd complex (Chu et al., 2013; Hirai et al., 2019; Janda et al., 2012). The Wnt-Fzd-Lrp complex inhibits the negative regulator destruction complex and stabilizes the Wnt signaling transducer β-catenin, which allows the transcription of genes regulating embryonic development and patterning (Bilić et al., 2007).

Wnt3 proteins, a subset of the Wnt family, are instrumental in the development of the nervous system, vascular system, limb formation and vertebrate axis formation (Anne et al., 2013; Bulfone et al., 1993; Clements et al., 2009; Garriock et al., 2007; Liu et al., 1999). In zebrafish, Wnt3 directs neural stem cell proliferation and differentiation, making it indispensable for brain development (Clements et al., 2009). Our group showed that in zebrafish embryos, Wnt3 associates with domains on the membrane (Azbazdar et al., 2019; Ng et al., 2016; Sezgin et al., 2017). Blocking the activity of Porc and thus reducing Wnt acylation by the drug C59, resulted in reduced domain confinement and defective brain development in zebrafish embryos (Ng et al., 2016; Teh et al., 2015). The understanding of the Wnt3 action mechanism in zebrafish brain development, therefore, requires identifying its source regions, determining its mode of transport, demarcating receiving target tissues and measuring Wnt3-receptor interactions.

In this study, we first mapped the source and target regions of Wnt3 in the zebrafish brain by comparing the expression of a transgenic line expressing functional Wnt3EGFP, with a reporter line expressing an inner plasma membrane targeting sequence tagged with mApple (PMTmApple). The expression in both lines are regulated by a 4 kb *wnt3* promoter that contains most of the regulatory elements and reports the spatiotemporal expression of *wnt3* (Teh et al., 2015). Wnt3EGFP spreads from where it is produced, while PMTmApple remains attached to the inner membrane leaflet of the producing cells. Hence, by analyzing the expression patterns of Wnt3EGFP and PMTmApple, we were able to identify the midbrain-hindbrain boundary (MHB), the brain midline (roof plate and floor plate) and the epithalamus as source regions of Wnt3, and the optic tectum (OT) and ventral regions of the cerebellum as distal target regions. Subsequently, we probed how Wnt3 is distributed from the source to the target regions of the zebrafish brain by measuring its *in vivo* dynamics using fluorescence correlation spectroscopy (FCS) and fluorescence recovery after photo-bleaching (FRAP). FCS is a single molecule sensitive technique that statistically analyzes the intensity fluctuations in a small observation volume (∼ femtoliter scale) to generate an autocorrelation function, from which the diffusion coefficient and the concentration of the fluorescent molecules in the observation volume are accurately evaluated (Enderlein et al., 2005; Kim et al., 2007; Krichevsky & Bonnet, 2002; Magde et al., 1974). FRAP, on the contrary, is an ensemble technique that measures the dynamics of the fluorescent molecules in a large region of interest (∼ micrometer scale) based on the recovery of the fluorescence intensity in an irreversibly photobleached region (Klonis et al., 2002; Koppel et al., 1976). FCS and FRAP both measure molecular mobilities and have been shown to provide consistent results (Macháň et al., 2016). However, as they access very different length scales, they can provide complementary information on local and global diffusion (Müller et al., 2012, 2013; Veerapathiran & Wohland, 2018).

Lastly, we monitored the *in vivo* interaction of Wnt3 with Fzd1, a potential target receptor, using fluorescence cross-correlation spectroscopy (FCCS) and calculated their binding affinity. In FCCS, the intensity fluctuations of two interacting molecules tagged with spectrally different fluorophores in an observation volume are cross-correlated, and their binding affinity *in vivo* is measured (Ries et al., 2009; Schwille et al., 1997; Shi et al., 2009). We observed that the coreceptor Lrp5 is essential for the interaction of Wnt3 with Fzd1. Our findings show that Wnt3 spreads from its source to target regions by extracellular diffusion governed by interactions with HSPG and its receptors.

## Results

### Identifying the Source and Target Regions for Wnt3

In order to identify the source and target regions of Wnt3, we used two transgenic lines: Tg(−4.0*wnt3:*Wnt3EGFP) and Tg(−4.0*wnt3:*PMTmApple). Tg(−4.0*wnt3:*Wnt3EGFP) is a functionally active Wnt3EGFP-expressing line, where Wnt3EGFP expression driven by 4 kb *wnt3* promoter (Teh et al., 2015). Tg(−4.0*wnt3:*PMTmApple) is a reporter line driven by the same 4 kb *wnt3* promoter, expressing PMTmApple. Since the 4 kb *wnt3* promoter contains most of the regulatory elements, Tg(−4.0*wnt3:*PMTmApple) is a faithful reporter of Wnt3 expression, which marks the plasma membrane of the Wnt3-producing cells. However, the localization of PMTmApple is restricted to its source cells, as it remains tethered to the inner leaflet of the plasma membrane. In contrast, the distribution pattern of Wnt3EGFP in Tg(−4.0*wnt3:*Wnt3EGFP) spans a broader range compared to PMTmApple in Tg(−4.0*wnt3:*PMTmApple), implying that Wnt3EGFP is transported from the source regions where it is produced to its distal target regions **(Figure 1)**. The overlap in the expression of the two lines, therefore, identifies the source regions, and the difference demarcates the distal target regions.

**Figure 1:**
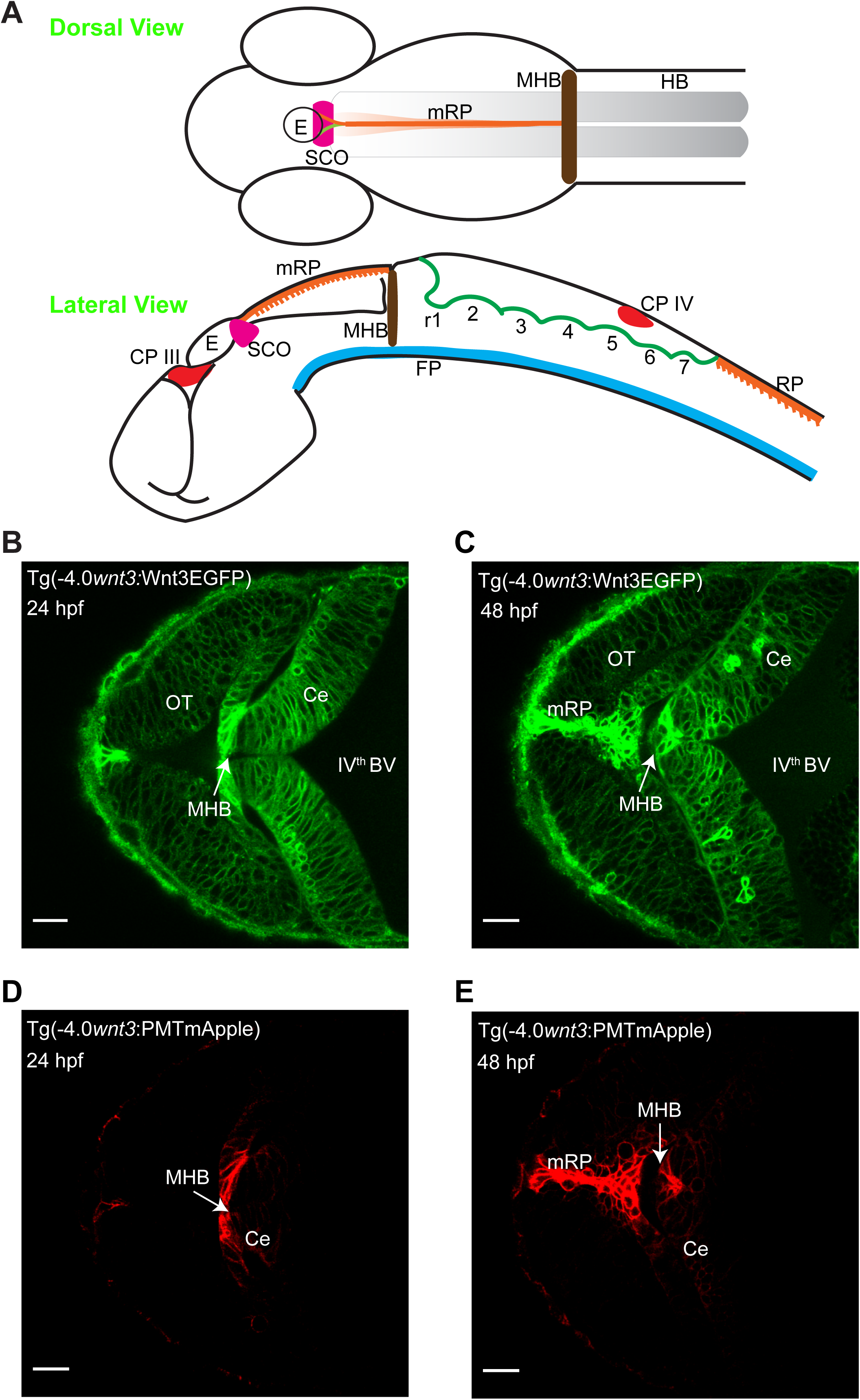
Spatiotemporal expression of *wnt3* promoter driven Wnt3EGFP and PMTmApple. **(A)** Schematic illustration of the brain of a zebrafish embryo (Dorsal view and Lateral view). Expression profile of Wnt3EGFP in Tg(−4.0*wnt3*:Wnt3EGFP) line at **(B)** 24 hpf and **(C)** 48hpf. Expression profile of PMTmApple in Tg(−4.0*wnt3*:PMTmApple) line at **(D)** 24 hpf and **(E)** 48hpf. BV, brain ventricle; Ce, cerebellum; CP, choroid plexus; E, epiphysis; FP, floor plate; HB, hindbrain; MHB, mid-brain-hindbrain boundary; mRP, midbrain roof plate; OT, optic tectum; r, rhombomere RP, roof plate (spinal cord); SCO, sub-commissural organ. Scale bar 30 μm.

The two transgenic lines were crossed [Tg(−4.0*wnt3:*Wnt3EGFP) × Tg(−4.0*wnt3:*PMTmApple)] and the expression of Wnt3EGFP and PMTmApple were sequentially recorded using a confocal microscope in their respective wavelength channels. Firstly, the obtained image stacks were segmented using an automatic threshold algorithm (Zhu et al., 2016), and the colocalization of each pixel was evaluated based on the intensity correlation analysis (ICA), the distance weight and intensity weight (Li et al., 2004; Zhu et al., 2016) to generate a pair of masks for the colocalized and non-colocalized pixels. Subsequently, color-coded heat maps were generated, indicating the contribution of each pixel to the overall colocalization at 24 and 48 hpf **(Video 1A and 1B)**. Finally, using the colocalized and non-colocalized masks, volumetric images were constructed to distinguish the source and target regions of Wnt3 respectively. At 24 hpf, the source regions were midbrain-hindbrain boundary (MHB), dorsal cerebellum (dCe), and epithalamus (Epi); whereas the distal target regions were ventral cerebellum (vCB) and optic tectum (OT) **(Figure 2 and Video 2)**. The source regions at 48 hpf were the midbrain roof plate (mRP), floor plate (FP), mid-brain-hindbrain boundary (MHB), dorsal cerebellum (dCe), epithalamus (Epi), and some parts of the dorso-lateral optic tectum (dOT); while the distal target regions were ventral cerebellum (vCe), and ventral optic tectum (vOT) **(Figure 3 and Video 3).** The mapped source regions agreed with the *in vivo* expression pattern of the *wnt3* gene as shown at mRNA level by whole mount *in situ* hybridization (**Supplementary Figure 1).** With the source and target regions defined, we next quantified the dynamics of Wnt3, and examined the mode of dispersal of Wnt3 from the source to the distal target regions.

**Figure 2:**
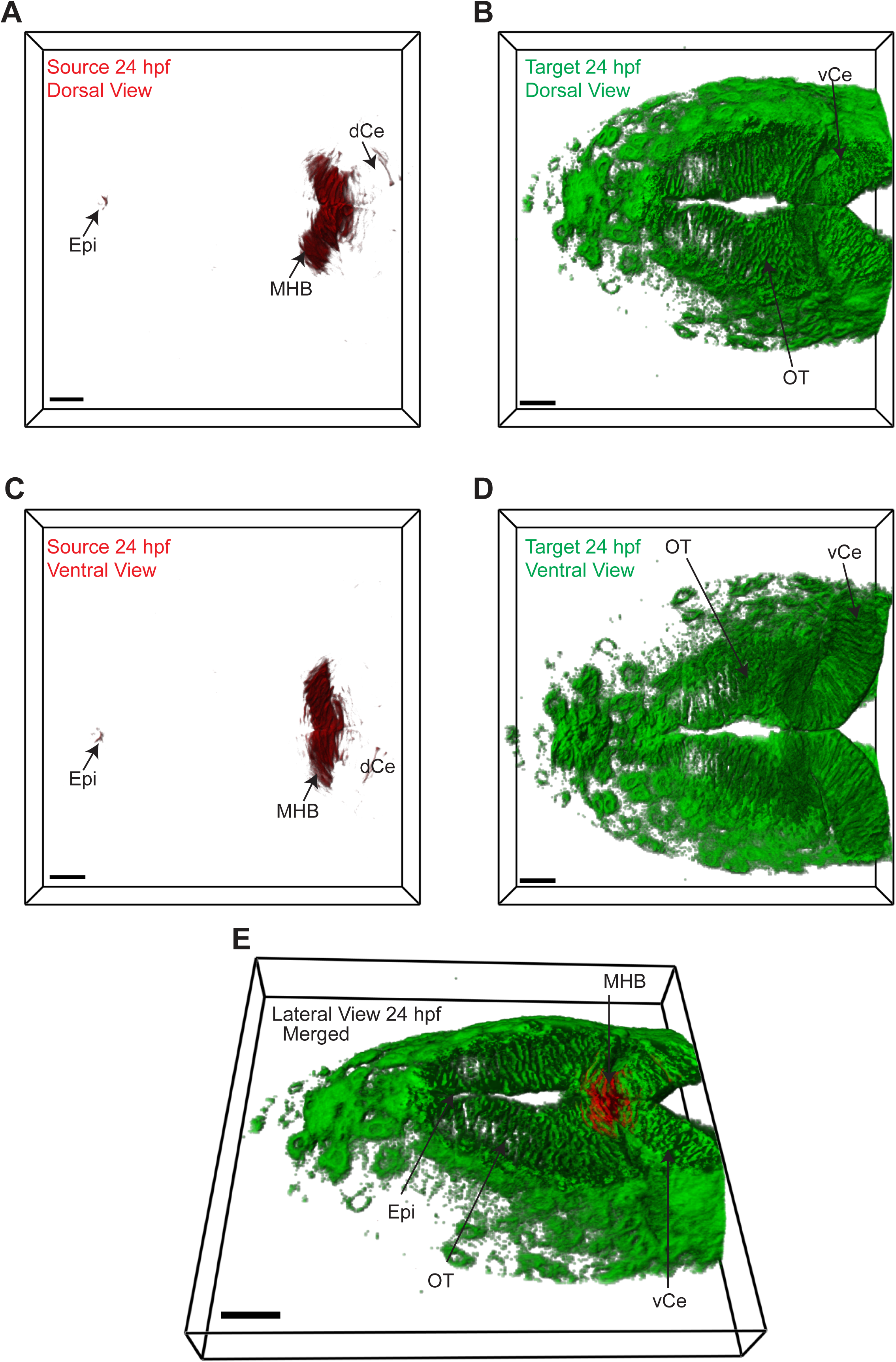
Identifying Wnt3 source and target regions at 24 hpf. 3D dorsal projection of Wnt3 **(A)** source regions at 24 hpf and **(B)** target regions at 24 hpf (Top view). 3D ventral projection of Wnt3 **(C)** source regions from at 24 hpf and **(D)** target regions at 24 hpf (Bottom view). **(E)** 3D projection of Wnt3 source and target regions from at 24 hpf (Lateral view). See Video 2 for a detailed view. dCe, dorsal regions of cerebellum; Epi, epithalamus; MHB, midbrain-hindbrain boundary; OT, optic tectum; vCe, ventral regions of cerebellum. Scale bar 30 μm.

**Figure 3:**
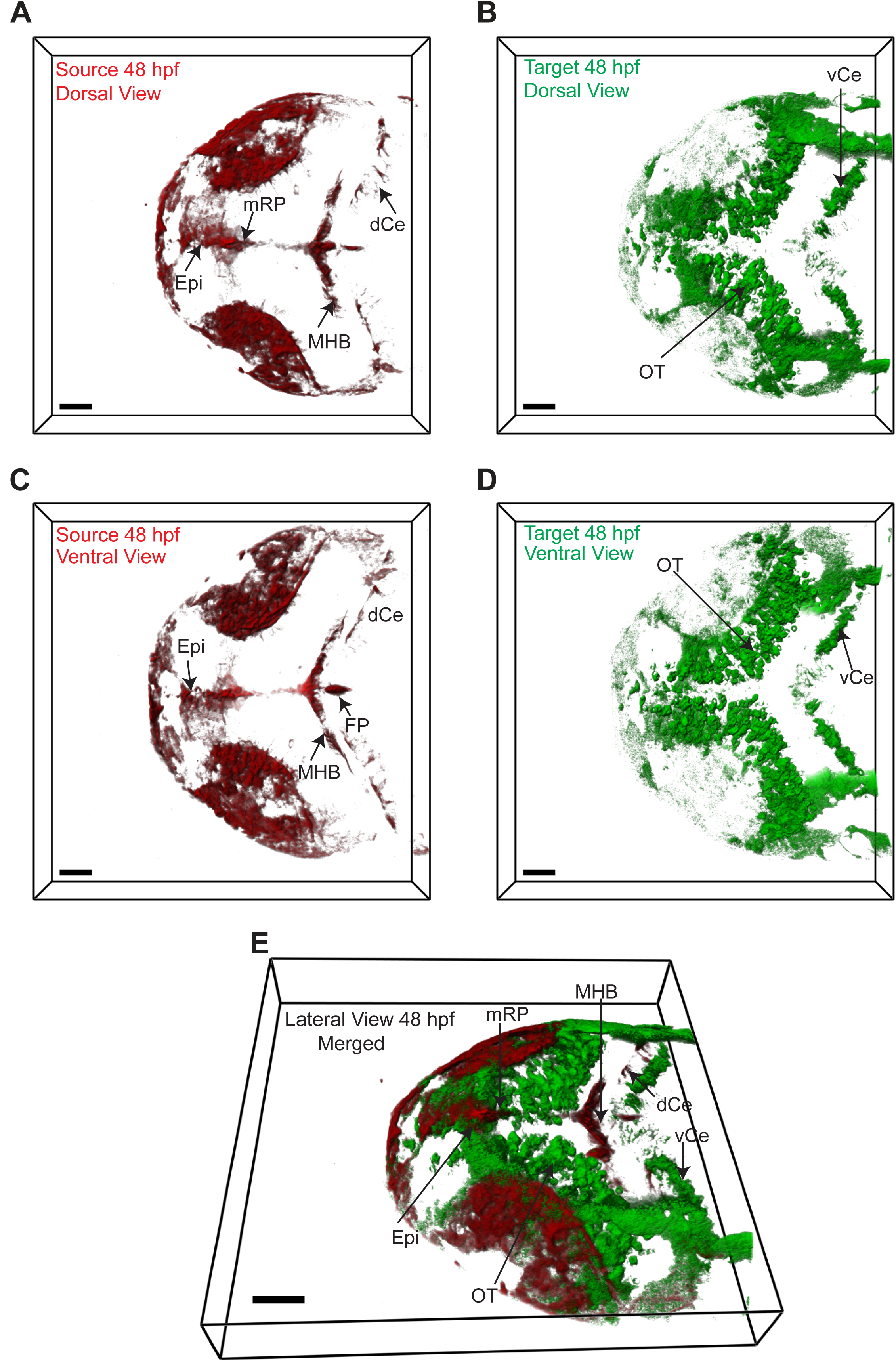
Identifying Wnt3 source and target regions at 48 hpf. 3D dorsal projection of Wnt3 **(A)** source regions at 48 hpf and **(B)** target regions at 48 hpf (Top view). 3D ventral projection of Wnt3 **(C)** source regions from at 48 hpf and **(D)** target regions at 48 hpf (Bottom view). **(E)** 3D projection of Wnt3 source and target regions from at 48 hpf (Lateral view). See Video 3 for detailed view. dCe, dorsal regions of cerebellum; Epi, epithalamus; FP, floor plate; MHB, midbrain-hindbrain boundary; mRP, midbrain roof plate; OT, optic tectum; vCe, ventral regions of cerebellum. Scale bar 40 μm.

### Characterizing the *in vivo* dynamics of Wnt3EGFP

At 24 and 48 hpf the cells are densely packed in the cerebellum, and optic tectum and there is no apparent extracellular space resolved within the limits of our microscopes (∼200 nm). It is thus not possible to determine from imaging alone whether Wnt3 is present in the interstitial spaces. We therefore use an indirect approach and measure the molecular mobility of Wnt3 at the borders between neighboring cells **(Figure 4 A, B)**. As diffusion coefficients on membranes and in aqueous solution differ by at least one order of magnitude if not more, they can be easily distinguished, and the presence of a freely diffusible species can be identified. For FCS measurements along the cell borders, we used a 2D-2particle-1triplet model (See Materials and Methods equation 7) as the fit model, as determined by the Bayes inference-based model selection (Sun et al., 2015; Teh et al., 2015). The fact that data can be fit with a 2D model most likely indicates that Wnt3 either diffuses on the membrane or in the narrow interstitial spaces that have a very small extent (< 200 nm) compared to the axial extent of the confocal volume (∼1 μm). We detected two diffusive components from these measurements: a slow component with a diffusion coefficient (D_slow_) of 0.6 ± 0.3 µm^2^/s and a fast component with a diffusion coefficient (D_fast_) of 27.6 ± 3.9 µm^2^/s **(Figure 4E).** The slow diffusive component was the dominant fraction (F_slow_ ∼ 0.6 ± 0.05) and represents the fraction of Wnt3 on the membrane. Note, however, that we cannot unambiguously assign the fast diffusion coefficient to Wnt3 in the interstitial spaces. The confocal volume for FCS measurements on the membrane also spans a portion of the intracellular cytosol. Hence, a fraction of Wnt3 within the cytosol could have contributed to the fast diffusion. Therefore, we tested whether the fast diffusion coefficient is susceptible to changes in the interstitial spaces, as discussed in the following section.

**Figure 4:**
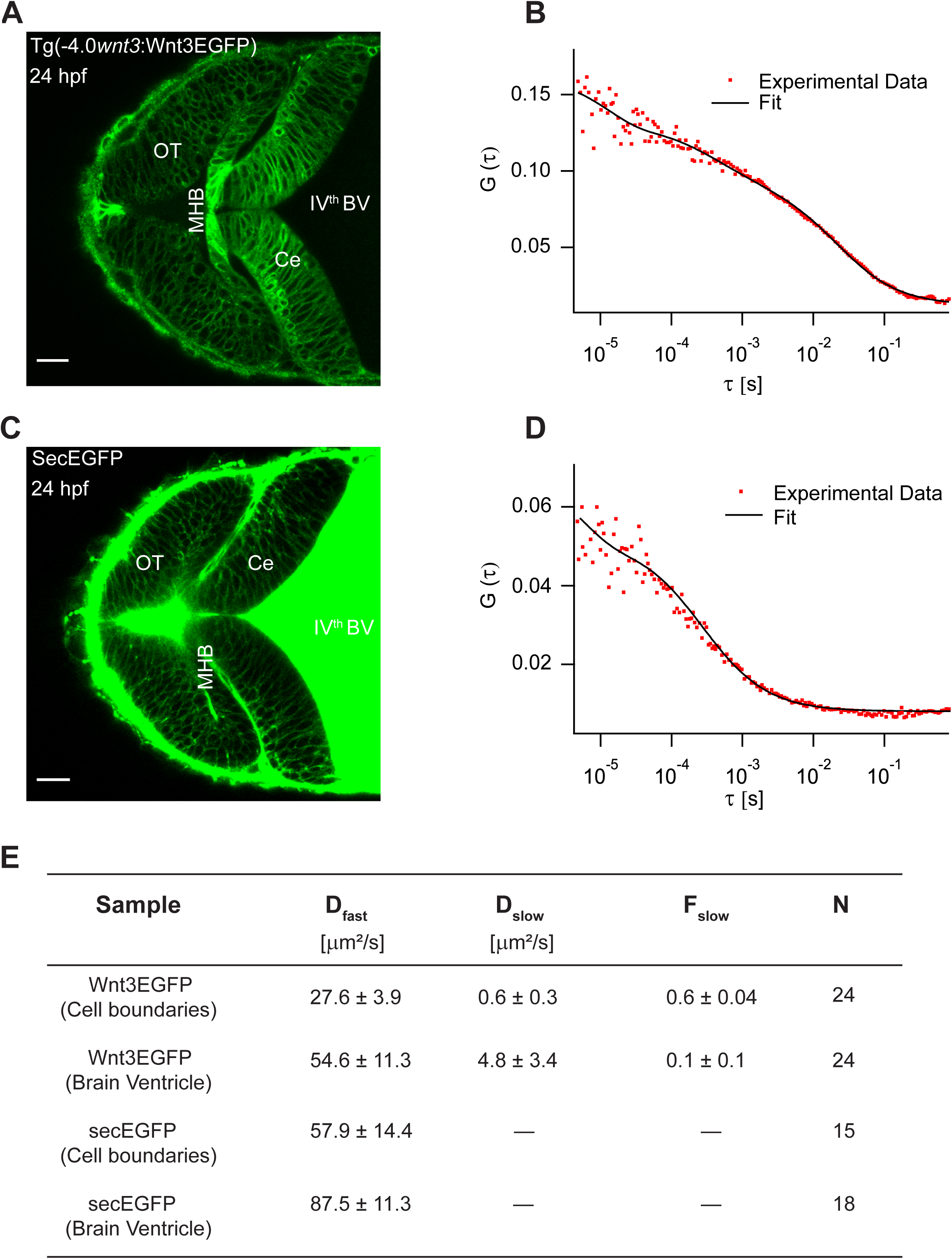
Determination of diffusion coefficients of Wnt3EGFP and secreted EGFP by FCS. **(A)** Expression of Wnt3EGFP in transgenic Tg(−4.0*wnt3*:Wnt3EGFP) line. **(B)** Representative autocorrelation function (dots) and fitting (line) of Wnt3EGFP measurement at cell boundary. **(C)** Expression of secEGFP in zebrafish brain. **(D)** Representative autocorrelation function (dots) and fitting (line) of secEGFP measurement at a cell boundary. **(E)** Table showing diffusion coefficients of the fast component (D_fast_), slow component (D_slow_) and the fraction of slow component (F_slow_) for Wnt3EGFP and secEGFP. Data are Mean ± SD; N = No of measurements. BV, brain ventricle; Ce, cerebellum; MHB, midbrain-hindbrain boundary; OT, optic tectum. Scale bar 30 µm.

### Wnt3 spreads extracellularly in the interstitial spaces

As Wnts are highly hydrophobic molecules, they tend to aggregate after being secreted into the extracellular milieu, which would limit them to autocrine and juxtacrine signaling (Fuerer et al., 2010). However, the expression of Wnt3EGFP in Tg(−4.0*wnt3:*Wnt3EGFP) was detected at a distance (∼ 50 – 150 µm) from the recognized source regions, implying long-range travel. Hence, we examined how Wnt3 spreads across the zebrafish brain, and whether it chooses the extracellular route. Since the cells are tightly packed at late stages (after 24 hpf) of the zebrafish embryo, we first verified the existence of the interstitial spaces at these late stages. We injected secreted EGFP (secEGFP), the secretory peptide of Fibroblast growth factor 8a (Fgf8a) tagged with EGFP, at the one-cell stage and imaged the zebrafish brain at 48 hpf. The secEGFP is targeted for extracellular secretion after their translation in the cytoplasm and a marker of the interstitial spaces. We observed the expression of secEGFP along the cell boundaries of the zebrafish brain and in the brain ventricles **(Figure 4C)**. When the dynamics for secEGFP was measured using FCS, we obtained a D of 57.9 ± 14.4 µm^2^/s along the cell boundaries **(Figure 4 D, E)**. As secEGFP does not bind to the cell membrane, this indicates its diffusion in the extracellular spaces, consistent with the fast diffusion coefficient measured (Müller et al., 2012, 2013).

As mentioned above, we were unable to unambiguously assign Wnt3 diffusion to its presence in interstitial spaces. Thus, we evaluated the effects of HSPG, a cell surface and extracellular matrix protein which should influence only molecules in interstitial spaces, on the dynamics of Wnt3. Since the interactions of Wnts with HSPG and the significance of HSPG in the activity of Wnts is well established (Fuerer et al., 2010; Kirkpatrick & Selleck, 2007; Mii et al., 2017), we treated the Tg(−4.0*wnt3:*Wnt3EGFP) embryos with heparinase to disrupt the HSPG and measured the dynamics of Wnt3EGFP. Injecting heparinase at the one-cell stage showed impaired gastrulation, so heparinase along with a high molecular weight fluorescent dextran (70,000 MW Dextran-TRITC) was co-injected in the brain ventricle of 48 hpf Wnt3EGFP expressing embryos. Since the presence of fluorescent dextran was detected along cell boundaries of the cerebellum and optic tectum, we inferred that heparinase (∼42 kDa) also diffused into the interstitial spaces from the brain ventricle **(Supplementary Figure 2)**. Confocal FCS measurements revealed that while the D_slow_ of Wnt3 for heparinase treated and untreated embryos remained the same, the D_fast_ for heparinase treated embryos was almost two-fold higher (D_fast_ = 43.4 ± 7.6 µm^2^/s) in comparison to the untreated embryos (D_fast_ = 24.7 ± 4.8 µm^2^/s) **(Table 1)**.

**Table 1:** Influence of Heparan Sulfate Proteoglycans on the dynamics of Wnt3EGFP, LynEGFP and secretedEGFP.

| *Sample* | *D<sub>fast</sub> [$\mu\text{m}^2/\text{s}$]* | *D<sub>slow</sub> [$\mu\text{m}^2/\text{s}$]* | *F<sub>slow</sub>* | *No. of measurements* |
| --- | --- | --- | --- | --- |
| **Wnt3EGFP** | $24.7 \pm 4.8$ | $0.6 \pm 0.3$ | $0.6 \pm 0.02$ | 47 |
| **Wnt3EGFP + Heparinase** | $43.4 \pm 7.6$ | $0.4 \pm 0.2$ | $0.6 \pm 0.04$ | 63 |
| **LynEGFP** | $39.1 \pm 11.2$ | $2.2 \pm 0.6$ | $0.7 \pm 0.04$ | 29 |
| **LynEGFP + Heparinase** | $40.1 \pm 9.5$ | $2.7 \pm 0.7$ | $0.6 \pm 0.04$ | 35 |
| **SecEGFP** | $59.4 \pm 9.4$ | - | - | 30 |
| **SecEGFP + Heparinase** | $56.9 \pm 9.7$ | - | - | 30 |

Data are Mean ± S.D. Unpaired two-tail t-test were performed between untreated and the corresponding heparinase treated embryos. For $D_{\text{fast}}$ , p-value was $< 0.0001$ for Wnt3EGFP and the difference was non-significant for LynEGFP and SecEGFP. The difference was non-significant for other parameters in all samples.

As controls, we measured the effects of heparinase treatment on the diffusion of secEGFP and LynEGFP (a non-functional membrane tethered tyrosine kinase). When secEGFP embryos were treated with heparinase, we observed no change in D compared to the untreated embryos **(Table 1)**. For LynEGFP in Tg(−8.0*cldnB*:LynEGFP), we obtained a slow component with a D_slow_ of 2.2 ± 0.6 µm^2^/s and a fast component with a D_fast_ of 39.1 ± 11.2 µm^2^/s. D_slow_ corresponds to the membrane diffusing component while D_fast_ represents a putative cytosolic fraction. When LynEGFP embryos were treated with heparinase, we did not observe any changes in D_fast_ or D_slow_, confirming that neither membrane nor cytosolic diffusion are influenced by HSPG disruption **(Table 1)**. This supports the hypothesis that Wnt3 is diffusing in the extracellular space and is regulated by interactions with HSPG.

To substantiate our results, we determined the global diffusion of Wnt3 using FRAP. As FRAP measures mobility over a range of several cell diameters, it is an ideal tool to investigate whether Wnt3 can diffuse extracellularly or by other much slower cellular mechanisms. We irreversibly photobleached a region of the zebrafish brain in Tg(−4.0*wnt3*:Wnt3EGFP) embryos, and observed the rate of recovery in the photobleached region. On analyzing the FRAP curve for Wnt3EGFP, two components with different recovery rates were obtained: a fast component with a recovery rate (τ_fast_) of 5-8 minutes and a slow component with a recovery rate τ_slow_ > 40 minutes. The mobile fraction (F_m_) of Wnt3EGFP evaluated from the FRAP curve was 0.3-0.4 with an effective diffusion coefficient (D_eff_) of 0.5 ± 0.2 µm^2^/s **(Figure 5)**. When FRAP was performed for secEGFP and PMTmApple expressing embryos in the same region of the zebrafish brain, secEGFP showed rapid recovery of 13 - 30 s (with F_m_ of 0.7 – 0.9 and D_eff_ of 13 ± 4 µm^2^/s) **(Supplementary Figure 3)**, while PMTmApple showed no recovery within the same measurement time **(Supplementary Figure 4)**. Although the source regions continuously produce PMTmApple, the generation of novel PMTmApple involves transcription, translation, and post-translational chromophore maturation (maturation time for mApple is ∼ 30 minutes). Since PMTmApple is tethered to the cell membrane, no recovery is observed after photobleaching of PMTmApple before 30 minutes. Hence, the recovery within 5-8 minutes for Wnt3EGFP points towards an extracellular distribution of Wnt3EGFP in the interstitial spaces of the developing zebrafish brain but with almost twenty-five-fold reduced D compared to secEGFP.

**Figure 5:**
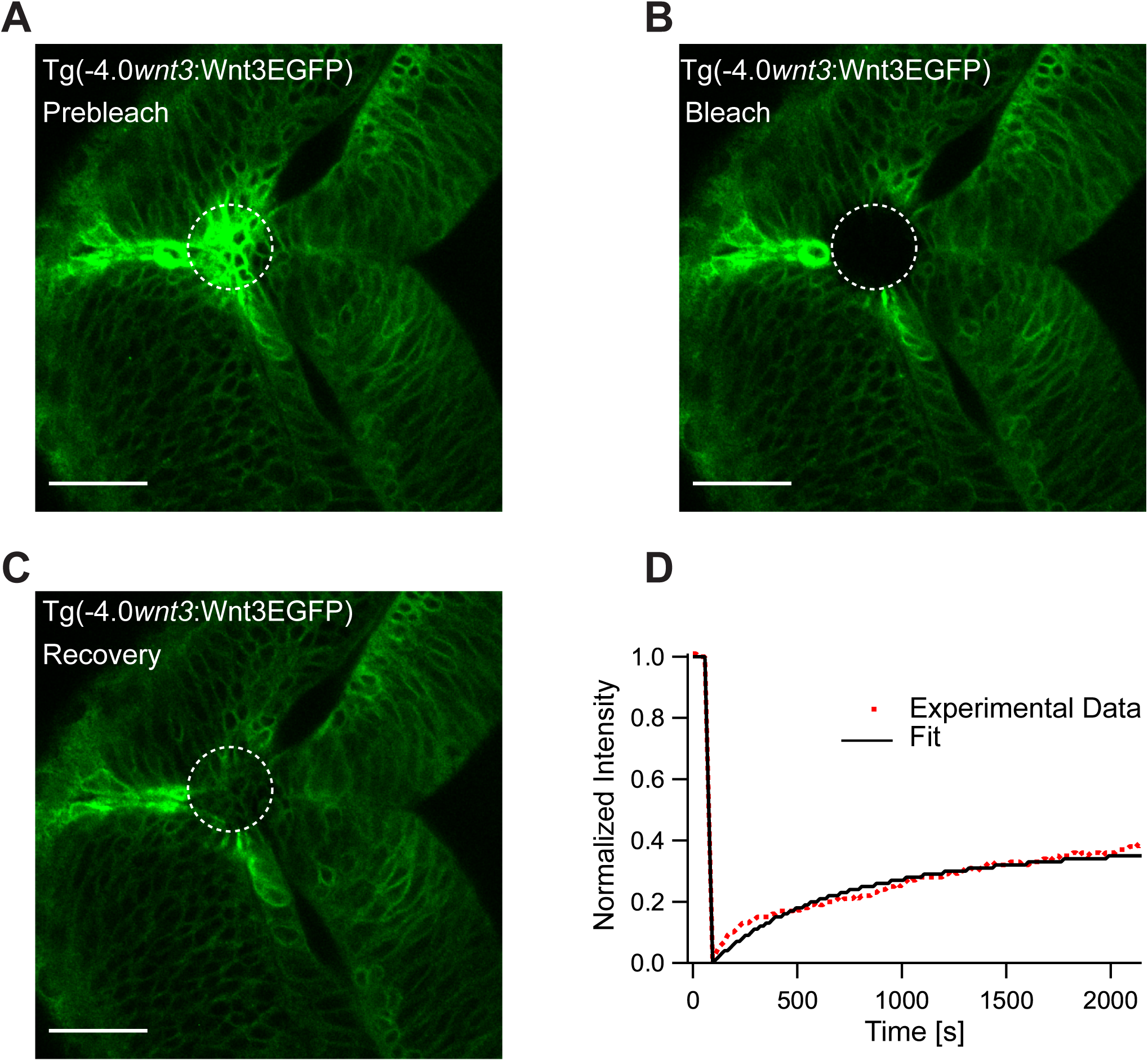
Representative fluorescence recovery of Wnt3EGFP after photobleaching. **(A)** Expression of Wnt3EGFP in Tg(−4.0*wnt3*:Wnt3EGFP) before photobleaching. **(B)** Photobleached region of Wnt3EGFP. **(C)** Recovery of fluorescence intensity in the bleached region due to diffusion of molecules from the neighboring unbleached regions. **(D)** Fluorescence recovery curve for Wnt3EGFP with a recovery time (τ_fast_) of ∼ 5 minutes and a mobile component fraction (F_m_) of ∼ 0.35. The average global diffusion coefficient (D_eff_) measured for Wnt3EGF P was 0.5 ± 0.2 μm^2^ /s (N=11). Scale bar 30 µm.

### The *in vivo* interactions of Wnt3 with Fzd1 receptor depends on the expression of *lrp5* coreceptor

Apart from the interactions of signaling molecules with the extracellular matrix proteins, the transient trapping of ligands by their receptors also shapes their distribution profile (Müller et al., 2013). For instance, the transient binding of Nodals to their receptor Acvr2b and co-receptor Oep (Lord et al., 2019; Wang et al., 2016), Hedgehog to the 12-transmembrane protein Dispatched (Callejo et al., 2011) and Wingless to the Fzd receptor (Baeg et al., 2004) influence their respective distributions and gradient kinetics. Hence, it is critical to evaluate the binding affinity of Wnt3 with its target receptors to understand its signaling range and action. Although the binding affinities for different Wnt ligands and Fzd receptors were quantified, they were limited to biochemical analysis on mammalian cell lines (Dijksterhuis et al., 2015). The dynamics and conformation of proteins might differ significantly *in vivo* (Lipinski & Hopkins, 2004), and quantitative analysis of Wnt-Fzd interactions in live organisms is still lacking. Since *in vitro* genetic and biochemical assays reported that Wnt3 interacts strongly with Fzd1 (Dijksterhuis et al., 2015), we investigated the *in vivo* Wnt3-Fzd1 interaction and measured its binding affinity. For this purpose, we generated a transgenic line Tg(−4.0*wnt3*:Fzd1mApple) expressing Fzd1mApple, crossed it with the Wnt3EGFP expressing line, and studied *in vivo* interactions using quasi-PIE FCCS **(Figure 6 A,B).** Quasi-PIE FCCS is an extension of FCCS, where the sample is simultaneously illuminated by a pulsed laser line and a continuous wave laser line of different wavelengths (Padilla-Parra et al., 2011; Yavas et al., 2016). This approach allows us to filter the background, spectral cross-talk, and detector after pulsing while computing the auto- and cross-correlation functions (Kapusta et al., 2012). When quasi-PIE FCCS measurements were performed in embryos expressing Wnt3EGFP and Fzd1mApple, we obtained cross-correlation between the two channels, indicating the *in vivo* interaction of Wnt3 with Fzd1 **(Figure 6C)**. As a positive control, we used embryos expressing PMT-mApple-mEGFP, and as negative control, we used embryos expressing Wnt3EGFP and PMTmApple by crossing their respective transgenic lines **(Supplementary Figure 5)**. The auto- and cross-correlations were then fitted with equation (7), and the binding affinity was measured according to equation (12) **(See Materials &Methods).** We obtained an apparent dissociation constant (K_d_) of 112 ± 15 nM indicating that Wnt3 binds strongly with Fzd1 *in vivo* **(Figure 6D)**. The measured *in vivo* K_d_ for Wnt3-Fzd1 is comparable with the *in vitro* K_d_ values reported for Wnts with Fzd1, which were in the range of 15 – 90 nM (Dijksterhuis et al., 2015).

**Figure 6:**
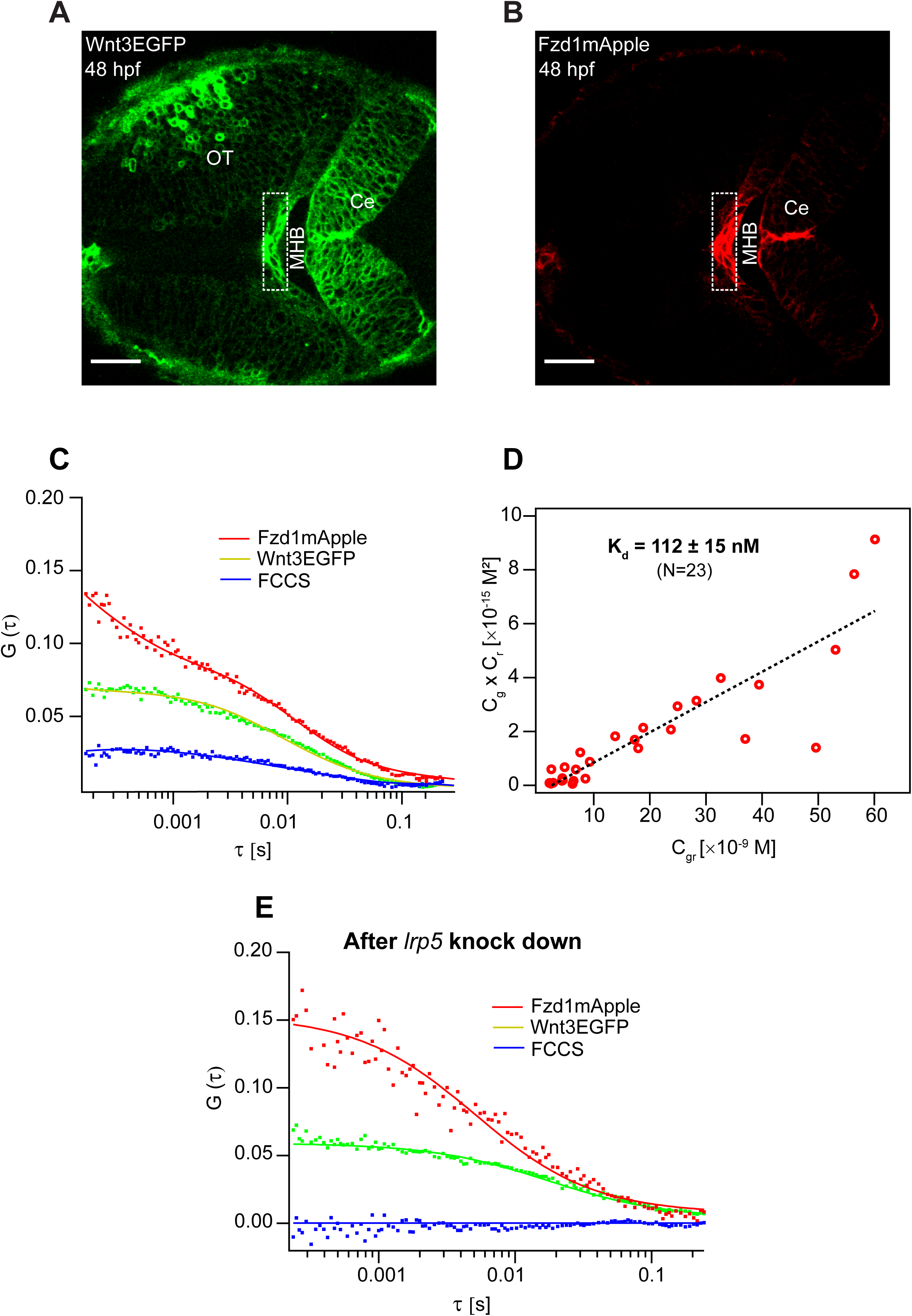
Determination of *in vivo* Wnt3-Fzd1 binding affinity by FCCS. Expression of **(A)** Wnt3EGFP and **(B)** Fzd1mApple in the double transgenic [Tg(−4.0*wnt3*:Wnt3EGFP) × Tg (−4.0*wnt3*:Fzd1mApple)]. **(C)** Representative auto- and cross-correlation functions (dots) and fittings (lines) of a Wnt3EGFP-Fzd1mApple measurement at the indicated region. The cross-correlation function indicates Wnt3 interacts with Fzd1 *in vivo*. **(D)** Determination of apparent dissociation constant (K_d_) for Wnt3EGFP-Fzd1mApple interaction *in vivo.* Cg, Cr, and Cgr represent the concentration of unbound Wnt3EGFP, unbound Fzd1mApple and bound Wnt3-Fzd1 molecules respectively. The estimated apparent K_d_ [K_d_ = (Cg * Cr) / Cgr] for Wnt3-Fzd1 *in vivo* is 112 ± 15 nM (N=23, R2 = 0.85). **(E)** Representative auto- and cross-correlation functions (dots) and fittings (lines) of a Wnt3EGFP-Fzd1mApple measurement after knocking down of the expression of *lrp5*. No cross-correlation indicates Wnt3-Fzd1 interaction is abolished after knockdown of *lrp5*. Scale bars 30 μm

Interestingly, Wnt3-Fzd1 interactions were only detected in the MHB and the dorsal cerebellum of the zebrafish brain at 48 hpf. No interactions were detected in the ventral cerebellum or optic tectum despite detecting Wnt3 and Fzd1 in these regions. Since the expression of the co-receptor *lrp5* corresponds to the specific regions where we detected interactions (Willems et al., 2015), we hypothesized that Lrp5 is necessary for the *in vivo* binding of Wnt3 to Fzd1. To test this, we knocked down the expression of *lrp5* using morpholinos (Mo) and checked for Wnt3-Fzd1 interactions in the MHB and dorsal cerebellum. We did not detect any cross-correlations after Mo treatment in these regions, whereas cross-correlations were obtained in the corresponding regions for untreated embryos (**Figure 6E).** When we performed FRAP experiments for the Mo-injected embryos, we obtained a faster recovery of ∼ 2 minutes for Wnt3EGFP in the photobleached regions with a D_eff_ of 3 ± 0.8 µm^2^/s **(Supplementary Figure 6)**. These results suggest that the co-receptor Lrp5 is essential for *in vivo* interaction of Wnt3 with Fzd1 and that this interaction influences Wnt3 diffusion.

## Discussion

Symmetry breaking and the development of an embryo into an organism requires a finely balanced but robust position-sensitive control of cell behavior and differentiation. This is achieved by signaling molecules that are expressed in well-defined source regions and distribute to target tissues where they are recognized by their cognate receptors. Wnts are a class of molecules that fulfil this function and are involved in cell division, cell migration, apoptosis, embryonic axis induction, cell fate determination, and maintenance of stem cell pluripotency (Clevers & Nusse, 2012; Logan & Nusse, 2004). Misregulation of this process leads to developmental defects and diseases, including cancer. In this work, we investigated the *in vivo* action mechanism of Wnt3, a member of this family that is involved in the proliferation and differentiation of neural cells, with particular attention to the differentiation of source and target regions, the mode of transport and the recognition of Wnt3 at the target site.

First, we analyzed the colocalization of Wnt3EGFP and PMTmApple expression in the double transgenic [Tg(−4.0*wnt3:*Wnt3EGFP) × Tg(−4.0*wnt3:*PMTmApple)] to map Wnt3 source and distal target regions at 24 hpf and 48 hpf. We categorized the MHB, midbrain roof plate, floor plate, epithalamus, and dorsal regions of the cerebellum as the source regions for Wnt3. Interestingly, earlier studies had documented these regions as the primary signaling centers that control the development of the central nervous system (CNS). The brain midline, comprising of the roof plate and floor plate, represent the signaling glia that acts as the source of several secreted signals involved in the neuronal specification (Chizhikov et al., 2006; Jessell TM, 2000; Kondrychyn et al., 2013). Chizhikov and Millen provided a comprehensive overview on how the roof plate governs the specification of the hindbrain, diencephalon, telencephalon and spinal cord by producing BMP and Wnt proteins (Chizhikov & Millen, 2005). Similarly, the importance of the MHB (also known as the isthmic organizer) in the morphogenesis of the zebrafish brain is also well studied (Gibbs et al., 2017; Raible & Brand, 2004; Wurst & Bally-Cuif, 2001). Our results, at a molecular level, corroborate these functional studies, which examine the role of these signaling centers in coordinating brain development by producing critical signaling molecules.

While whole-mount *in situ* hybridization (WISH) is useful for visualizing the spatial gene expression patterns on fixed embryos at the level of mRNA, it does not provide information regarding the distribution of signaling proteins in live samples. Our approach based on the analysis of the distribution of proteins *in vivo* enabled us to validate not only the source regions, but also identify the ventral regions of the cerebellum and optic tectum as the target regions to where Wnt3 is transported. However, the whole list of Wnt3 target sites could be longer. Recently, it was shown that Wnt5A transported in the cerebrospinal fluid regulates the development of the hindbrain (Kaiser et al., 2019). Since previously we also detected Wnt3 diffusing in the brain ventricles (Teh et al., 2015), further detailed investigation is required to detect additional less obvious target sites. Nevertheless, the characterization of Wnt3 source and target regions of this work clearly indicates the presence of discrete Wnt3 producing- and receiving-cells in the developing brain of zebrafish embryos.

Second, we investigated the transport mechanism of Wnt3 in the zebrafish brain. The transport mechanism not only influences signaling and function, but is of particular interest for Wnts as it is not clear how they can distribute over long distances despite their hydrophobic nature. Using FCS, we first quantified the *in vivo* dynamics of Wnt3EGFP along the cell boundaries and in the brain ventricle. In the brain ventricle we found two different diffusing components of 54.6 ± 11.3 µm^2^/s and a slow component with D_slow_ of 4.8 ± 3.4 µm^2^/s **(Figure 4E).** The first component is similar to secEGFP and is consistent with freely diffusing Wnt3EGFP, or at best Wnt3EGFP in a very small complex, e.g. with a shuttling protein that hides the hydrophobic Wnt3 moiety and prevents Wnt3 aggregation. The second component is much slower and hints at Wnt3EGFP associated with larger protein or lipid complexes and would be consistent with either exosomes or protein transport complexes. It will be interesting to address the exact nature of the aggregation and/or complexation state of Wnt3 in future studies. At the cell boundaries, we found two diffusive components for all Wnt3EGFP measurements, one that is consistent with membrane diffusion (D_slow_ = 0.6 ± 0.3 µm^2^/s) and another component (D_fast_ = 27.6 ± 3.9 µm^2^/s) too fast to be attributed to diffusion within a lipid bilayer and much closer to the diffusion coefficient seen for secEGFP (D = 57.9 ± 14.4 µm^2^/s). Due to resolution limitations of FCS, we could not unambiguously attribute this component to secreted Wnt3EGFP, as cytosolic Wnt3EGFP could also contribute to the fast diffusing component. Since Wnt3 has been shown to interact with HSPG (Fuerer et al., 2010; Kirkpatrick & Selleck, 2007; Mii et al., 2017), we disrupted HSPG by heparinase injection, which should influence only extracellular Wnt3 but not a putative cytosolic component. In subsequent measurements, D_fast_ for Wnt3EGFP increased to 43.4 ± 7.6 µm^2^/s upon heparinase treatment indicating that Wnt3 spreads by extracellular diffusion.

FRAP experiments conducted at 48 hpf as target region and multiple cell diameters removed from the source region corroborate these results. Fluorescence recovery took place within 7 minutes indicating transport over long distances. However, the estimated effective diffusion coefficient of Wnt3EGFP was only 0.5 ± 0.2 µm^2^/s, a factor ∼50-100 lower than the diffusion coefficient in the interstitial spaces measured by FCS (27.6 ± 3.9 µm^2^/s). This is in stark contrast to the secEGFP global diffusion coefficient which was estimated to be 13 ± 4 µm^2^/s, and was reduced by only about a factor 3-5 compared to FCS measurements of the same molecule (57.9 ± 14.4 µm^2^/s). This smaller reduction in the global versus the local diffusion coefficient for secEGFP, as measured by FCS and FRAP respectively, could be an effect of tortuosity (Müller et al., 2013). However, the much larger reduction of the global diffusion coefficient for Wnt3EGFP calls for a different explanation, possibly including transient binding to its receptors and HSPG (Müller et al., 2013). Subsequent experiments showed that HSPG disruption by heparinase increased Wnt3EGFP diffusion by a factor ∼2 (FCS), and *lrp5* knockdown increased the Wnt3EGFP global diffusion coefficient by a factor ∼5-6. Overall this accounts for a reduction of global Wnt3EGFP diffusion by at least a factor 30-60, consistent with the 50-100 fold reduction seen by the comparison of shortrange (FCS) and long-range (FRAP) diffusion of Wnt3 in native conditions. Hence, our FCS and FRAP results collectively implicate the extracellular diffusion of Wnt3 mediated by HSPG and receptor binding to accomplish long-range dispersal in the developing zebrafish brain.

However, it must be noted that Wnt3 might additionally assume other modes of spreading. It is possible that carrier proteins or exosomes also shuttle Wnt3 in the zebrafish brain as would be consistent with the second slow component of Wnt3EGFP diffusion found in the brain ventricle. Moreover, HSPG may also assist in the transfer of Wnt bearing exosomes or lipoproteins by acting as their binding sites. A study demonstrated how HSPG guides the clearance of very low-density lipoprotein (VLDL) by forming a complex with Lrp (Wilsie & Orlando, 2003). Correspondingly, Eugster et al., explained how the interaction of the *Drosophila* lipoprotein with HSPG might influence the long-range signaling of Hedgehog in *Drosophila* (Eugster et al., 2007). On the same note, it was also determined how the functional activity of exosomes and vesicles is dependent on HSPG (Christianson & Belting, 2014). Thus, a detailed investigation is required to confirm if HSPG aids the transport of Wnt3 packaged in exosomes or lipoproteins in the zebrafish brain. Nevertheless, our findings illustrate how HSPG moderates the long-range extracellular spreading, and by extension the function, of Wnt3 in the zebrafish brain.

Once Wnt ligands reach the target cells, the next question is how they interact with their target receptors. As we had established that it is highly unlikely for Wnts to diffuse in the interstitial spaces freely, they must be released from their chaperones or HSPG in order to interact with their receptors. One possible hand-off mechanism is the competitive binding of Wnts to their target receptors with a higher binding affinity (Naschberger et al., 2017; Wilson, 2017). Furthermore, the binding affinity of the Wnt-receptor complex also modulates their range and magnitude *in vivo*. Hence, we measured the *in vivo* binding affinity for Wnt3 with a potential target receptor, Fzd1 using quasi-PIE FCCS. We obtained an apparent K_d_ of 112 ± 15 nM *in vivo*, implying a strong interaction. However, the actual K_d_ might be even lower as the concentration of the endogenous proteins, and the photophysics of the fluorophore affects the estimated K_d_ (Foo et al., 2012). Nonetheless, it is an estimate of the native *in vivo* Wnt3-Fzd1 binding in their physiological condition, which is consistent with results of *in vitro* experiments (Dijksterhuis et al., 2015). Interestingly, we also noticed that the interaction of Wnt3 with Fzd1 was dependent on the expression of the coreceptor Lrp5. We did not detect any cross-correlations when the expression of *lrp5* was knocked down and the D_eff_ for Wnt3EGFP increased by a factor ∼ 3-5. From this result, it appears that LRP5 is an essential component in facilitating the interaction of Wnt3 with Fzd1 with a significant influence on the diffusion coefficient and the long-range spreading of Wnt3. Hence, it is of interest to measure the K_d_ for Wnt3-LRP5 in the future and verify if the co-receptor is involved in the hand-off of Wnt from the carrier proteins and HSPG to its receptor. Note that Fzd1mApple expression in our transgenic line was driven by a 4 kb Wnt3 promoter that mimicked the regular expression of Wnt3. While useful methodologically to measure auto- and juxtacrine interactions of Wnt3-Fzd1, additional work is needed in measuring the *in vivo* binding affinities for Wnt3 with Fzd receptors expressed under the control of their native promoters.

In conclusion, our results show the presence of distinct Wnt3 source and target regions in the developing zebrafish brain, and that Wnt3 is distributed from its source to target by extracellular diffusion. We observed that the diffusion of Wnt3 is retarded by a factor 3-5 due to tortuosity, a factor 5-6 due to receptor binding and a factor ∼2 due to HSPG, thus leading to a total reduction of a factor 30-60 when comparing Wnt3EGFP short-range (∼ 28 µm^2^/s as measured by FCS) to long-range diffusion (∼ 0.5 µm^2^/s. as measured by FRAP). This indicates that the major part if not all the reduction seen for long-range compared to short-range diffusion of Wnt3 is explainable by tortuosity, receptor binding and interactions with HSPG present in the interstitial spaces.

Finally, we demonstrated that the co-receptor Lrp5 drives the *in vivo* interaction of Wnt3 with Fzd1, and quantitatively determined their affinity. This demonstrates that the presence of proteins alone, be it signaling molecules or receptors, as determined by fluorescence microscopy does not report on the actual signaling but it is necessary to measure interactions or downstream signaling to differentiate the concentration from the functional distribution of signaling molecules. Overall, our findings provide a general outline of Wnt3 signaling in the zebrafish brain from expression and transport to target binding, which set a starting point for the quantitative investigation of the Wnt3 interaction network during zebrafish brain development.

## Acknowledgements

We thank the NUS Centre for BioImaging Sciences, SingaScope, and the Institute of Molecular and Cell Biology for providing microscope facility support and providing zebrafish care. TW acknowledges funding by the Singapore Ministry of Education (MOE2016-T3-1-005). SV is supported by a NUS Research Scholarship. We thank David Piston for providing the construct of mApple.

## Author Contributions

SV and TW designed the experiments, analyzed and interpreted the results. SV performed the imaging, FCS and FRAP experiments. CT, VK and IK designed and generated the transgenic zebrafish lines. SZ performed the colocalization analysis. SV and TW wrote the manuscript and TW, VK, CT, SZ and PTM contributed in manuscript revision.

## Materials and Methods

### Fluorescence correlation spectroscopy

The molecular movement of fluorescently labeled molecules will cause fluorescence fluctuations during their entry and exit in a small open observation volume. These fluctuations contain the information about the dynamics of these molecules. In confocal FCS the confocal volume of the microscope setup defines the observation volume. The measured intensity trace is autocorrelated to extract the average concentrations and diffusion coefficients of the molecule in the sample. The autocorrelation function (ACF), G (τ), is given by

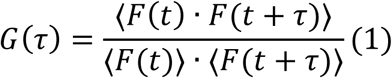

Where F(t) is the fluorescence intensity at time t, τ is the lag time and ⟨…⟩ represents time average. For a Gaussian illumination profile, G(τ) for a three dimensional free diffusion process with a single component and triplet state can be written as

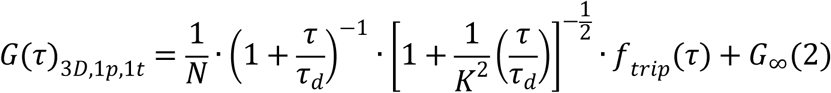

Here, *N* is the mean number of molecules in the observation volume and is inversely proportional to the amplitude of the ACF *G(0)* ; τ_d_ is the diffusion time of the molecule; *G*_∞_ is the convergence at long lag times; *K* is the structure factor which denotes the shape of the confocal volume

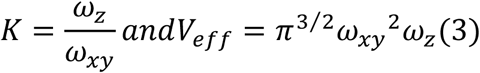

where *ω*_*z*_ and *ω*_*xy*_ are the 1/e^2^ radii of the PSF in the axial and radial direction; and *f*_*trip*_ *(τ)* is the triplet function which accounts for the fraction of particles in the triplet state *(F*_*trip*_) with a triplet relaxation time of *τ*_*trip*_, and it is represented as

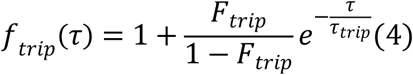

If two diffusing components are present, then the correlation function for two component 3D diffusion process *G (τ)*_*3D,2p,1t*_ is

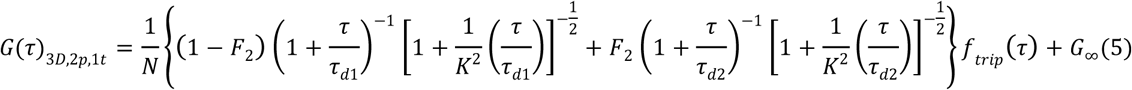

Where *F*_*2*_ is the fraction of the second component. For a 2D diffusion process such as on a membrane, the fitting equations (2) and (5) become

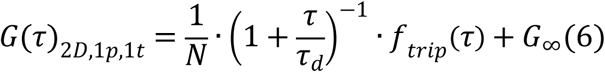

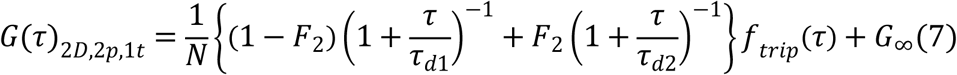

For FCS measurements, the system was first calibrated with Atto 488 dye for 488 nm and 485 nm laser lines and Atto 565 for 543 nm laser line. The known diffusion coefficient for the dye was 400 µm^2^/s at room temperature. The obtained correlation function was fit using equation (2) and the free fit parameters were N, τ, τ_trip_, F_trip_ and G_∞._ The K value and V_eff_ were calculated using equation (3). The samples were dechorionated, anesthetized by Tricaine and mounted in 1% low melt agarose in a No. 1.5 glass bottom MatTek petri dishes. The acquisition time for the measurements was 60 s and all measurements were performed at room temperature. For FCS measurements along the cell borders in Wnt3EGFP, LynEGFP, PMTmApple and Fzd1mApple expressing embryos, we used 2D-2particle-1triplet model (equation 7), and 2D-1particle-1triplet model (equation 2) in secEGFP expressing embryos. The measurements for Wnt3EGFP in the brain ventricle were fit using 3D-2particle-1triplet model (equation 5) and for secEGFP using 3D-1particle-triplet model (equation 2). The fit models were determined the Bayes inference-based model selection (Sun et al., 2015).

### Quasi PIE Fluorescence cross-correlation spectroscopy

FCCS is a valuable tool to study biomolecular interactions in live samples. When two interacting molecules tagged with spectrally different fluorophores transit through the observation volume, the intensity fluctuations from the two channels can be cross-correlated to obtain the cross-correlation function G_x_ (τ) given by:

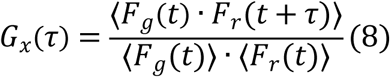

Where F_g_ and F_r_ are the fluorescence intensity in the green and red channel respectively.

For our FCCS measurements to detect Wnt3-Fzd1 interactions, we used an interleaved pulsed 485 nm laser line and a continuous wave 543 nm laser line to obtain the auto- and cross-correlation functions. This allowed us to apply statistical filtering (Kapusta et al., 2012) which helped in eliminating spectral cross-talk, background signal and detector after-pulsing based on fluorescence lifetime correlation spectroscopy (FLCS) as detailed in Parra et al., 2011 (Padilla-Parra et al., 2011). This is called quasi-PIE FCCS (Yavas et al., 2016).

Taking into account the background and spectral cross-talk, the amplitude of the ACF in the green channel G_G_(0), red channel G_R_(0), and the amplitude of the CCF G_x_(0) can be written as:

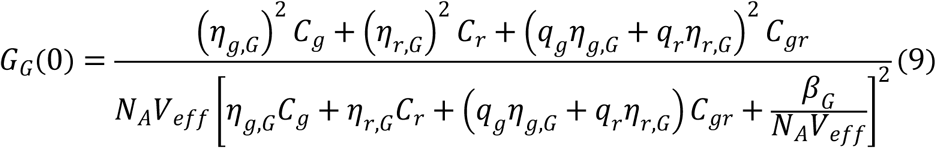

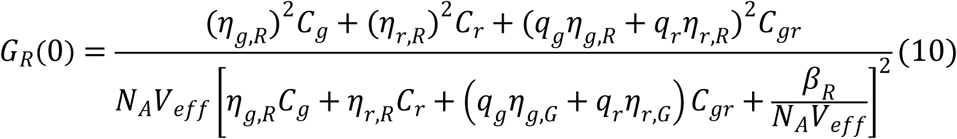

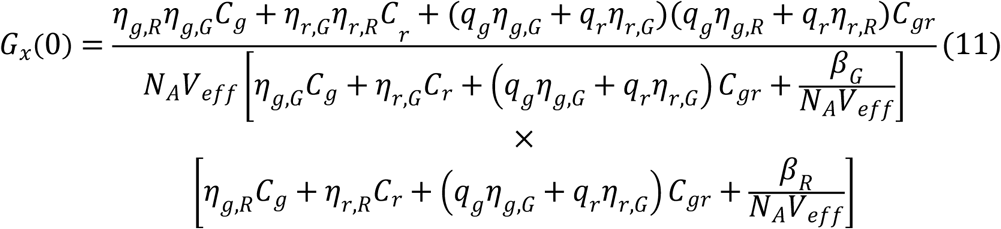

Here *η*_*g,G*_ and *η*_*r,R*_ represent the mean counts per particle per second (cps) for EGFP in the green and mApple in the red channel respectively. For our samples we obtained a *η*_*g,G*_ of ∼ 1900 Hz and *η*_*r,R*_ of ∼ 1400 Hz. *β*_*G*_ and *β*_*R*_ represent the count rates of background collected in the green and red channel respectively. *β*_*R*_ measured from blank WT embryo was ∼ 400 Hz while FLCS correction eliminated the background in the green channel (*β*_*G*_ = 0). N_A_ is the Avogadro’s number and V_eff_ represents the effective confocal volume from calibration.*η*_*r,G*_ and *η*_*g,R*_ denotes the cross-talk in the green and red channel respectively, which was efficiently eliminated by quasi PIE FCCS (*η*_*r,G*_ and *η*_*g,R*_ = 0). *q*_*g*_ and *q*_*r*_ are the correction factors due to FRET and quenching. Since the cps for Wnt3EFP and Fzd1mApple in their respective transgenics were same as that in double transgenic line, *q*_*g*_ and *q*_*r*_ were set to 1. Equations 9, 10 and 11 were solved for C_g_, C_r_ and C_gr_, which denote the concentration of the free green, free red and bound molecules in the observation volume respectively. Using C_g_, C_r_ and C_gr_ the dissociation constant (K_d_) for the interaction which can be determined using equation 12:

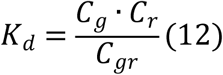

### Confocal microscope setup

An Olympus FV 1200 laser scanning confocal microscope (IX83; Olympus, Japan) integrated with a PicoQuant time resolved LSM upgrade kit (Microtime 200; GmbH, Germany) was used in this work. The sample was illuminated using a 488 nm laser beam (for EGFP) and 543 nm laser beam (for mApple) which was reflected to the back focal plane of an Olympus UPLSAPO 60X/1.2 NA water immersion objective. For all the experiments, the intensity of the laser before the objective was 20 µW. The emitted signal passes through a 120 µm pinhole before being filtered by an Olympus 510/23 emission filter (for EGFP) and Olympus 605/55 emission filter (for mApple) and eventually directed to a PMT detector for imaging. For FCS measurements, 510/23 emission filter (Semrock, USA) and 615DF45 filter (Semrock, USA) were used, and the filtered emissions were recorded using a single photon sensitive avalanche photodiodes (SAPD) (SPCM-AQR-14; PerkinElmer). The recorded signal then processed using SymPhoTime 64 (PicoQuant, Germany) to compute the autocorrelation function. For FCCS measurements, the sample was simultaneously illuminated with a pulsed 485 nm laser (LDH-D-C-488; PicoQuant) operated at 20 MHz repetition rate and a continuous 543 nm laser. The emission was separated using 560 DCLP dichroic mirror and directed to the 510/23 emission filter (Semrock, USA) and the 615DF45 filter (Semrock, USA). The signal recorded by SAPD was then analyzed by Synphotime to generate the auto- and cross-correlations.

### Colocalization analysis

For the colocalization analysis of Wnt3EGFP and PMTmApple, confocal z-stacks of step size 0.5 µm were obtained with identical acquisition settings. An automatic threshold algorithm detailed in Zhu et al., 2016 (Zhu et al., 2016)was implemented to segment the data. The algorithm uses the correlation quotient to select an optimal threshold for segmentation as described in (Li et al., 2004). Following the segmentation, the colocalization for each pixel was calculated based on intensity correlation analysis (ICA), the distance weight and intensity weight (Li et al., 2004; Zhu et al., 2016). Finally, a pair of masks for the colocalized and non-colocalized pixels were generated. The colocalized pixels and non-colocalized pixels were used to construct 3D images of the source and target regions respectively. The 3D images were built using ‘3D View’ module Imaris 9.5.0. The display setting was set to white background color and the 3D reconstructed images were represented in ‘Normal Shading’ mode for improving contrast in Figures 2, 3 and Videos 2, 3.

### Fluorescence recovery after photo bleaching (FRAP)

FRAP measurements were performed on an Olympus FV3000 laser scanning microscope. The mounted samples were imaged with a UPLSAPO 60X/1.2 NA water immersion objective using a 488 nm diode laser (for Wnt3EGFP and secEGFP) or a 561 nm diode laser (for PMTmApple). A DM 405/488/561/640 dichroic mirror separated the excitation and emission beams. The signal from the sample, after passing through the dichroic mirror was filtered by a BP 510-550 emission band pass filter for the 488 nm laser beam, and by a BP 575-625 emission band pass filter for the 561 nm laser beam. The pinhole size was adjusted to 1 AU. For FRAP, 5 pre-bleach frames were obtained before irreversibly photo bleaching a circular region of interest (ROI) for 30 seconds. The fluorescence intensity recovery in the photobleached region was recorded for 30 minutes. The images were then analyzed using the FRAP module in the Olympus CellSens software. A reference region on the sample but outside the ROI was selected to correct for photo bleaching, and another reference region outside the sample was selected for background correction. The software then plotted a FRAP recovery curve for the ROI, fitted the FRAP curve with a double exponential fit to obtain the recovery time for the fast (τ_fast_) and the slow (τ_slow_) component. The diffusion coefficient (D_eff_) was calculated using the Soumpasis equation (eq 13) for 2D circular bleaching of radius r (Kang et al., 2015; Koppel et al., 1976). However, it must be noted that this is an apparent estimate of D_eff_ as the distribution of fluorophores is not homogeneous, and we assume there is no diffusion during photo bleaching process.

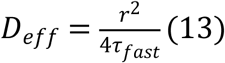

### Generation of transgenic lines and zebrafish maintenance

To generate Tg(−4.0*wnt3*:PMTmApple) transgenic zebrafish, the 45 bp plasma membrane targeting-sequence (PMT) (ATGGGCTGCTTCTTCAGCAAGCGGCGGAAGGCCGACAAGGA-GAGC) was cloned upstream and in-frame with mApple to generate PMTmApple open reading frame (ORF). The DNA fragment was subcloned into the 4-kbWnt3EGFP-miniTol2 recombinant plasmid (Teh et al., 2015) using Gibson assembly by replacing the Wnt3EGFP ORF with PMTmApple to give 4-kbPMT-mApple-miniTol2 recombinant plasmid.

To generate Tg(−4.0*wnt3*:Fzd1mApple), zebrafish *fzd1* ORF (1617 bp ; ENSDARG00000106062) was amplified by RT-PCR and subcloned into pGemTeasy. The Fzd1mApple DNA fragment was constructed by removing the Fzd1 stop codon and inserting in-frame (GGGS)2 linker sequence (GGAGGAGGATCAGGAGGAGGATCA) tagged with mApple to Fzd1 C terminal by Gibson assembly. This DNA fragment was then subcloned into the 4-kbWnt3EGFP-miniTol2 recombinant plasmid using Gibson assembly by replacing the Wnt3EGFP ORF with Fzd1mApple to give 4-kbFzd1mApple-miniTol2 recombinant plasmid.

Stable *wnt3* promoter-driven transgenic lines were generated as stated (Balciunas et al., 2006) by co-injection of transposase mRNA and 4-kbPMT-mApple-miniTol2 recombinant plasmid; co-injection of transposase mRNA and 4-kbFzd1mApple-miniTol2 recombinant plasmid, to generate Tg(−4.0*wnt3*:PMTmApple) and Tg(−4.0*wnt3*:Fzd1mApple) transgenic lines respectively.

Additional transgenic lines used are Tg(−8.0*cldn*B:lynEGFP) for *in vivo* imaging of membrane-tethered EGFP expression in the cerebellum (Haas & Gilmour, 2006). Wnt3EGFP expression in the brain was imaged using Tg(−4.0*wnt3*:Wnt3EGFP)F2 (Teh et al., 2015).

Transgenic adult zebrafish and embryos were obtained from zebrafish facilities in the Institute of Molecular and Cell Biology (Singapore) and National University of Singapore. The Institutional Animal Care and Use Committee (IACUC) in Biological Resource Center (BRC), A*STAR, Singapore (IACUC #161105) and the National University of Singapore (IACUC# BR18-1023) have approved the entire study. Spawned transgenic embryos were staged as described (Kimmel et al., 1995). Embryos older than 30 hpf were treated with 1-phenyl-2-thiourea at 18 hpf to prevent formation of melanin.

### Morpholino injection

The injected dose of *lrp5* splice-blocking Morpholinos (MOs; Gene Tools, Corvalis, USA) lrp5MoUp (AGCTGCTCTTACAGTTTGTAGAGAG) targeting the Exon2-Intron2 splice junction and lrp5MoDown (CCTCCTTCATAGCTGCAAAAACAAG) targeting the Intron2-Exon3 splice junction were conducted in accordance to published research (Willems et al., 2015).

### Heparinase injection into the zebrafish brain ventricle

Heparinase I from Flavobacterium heparinum (Merck) was dissolved in PBS to 1U/μl and stored as frozen aliquots. For microinjection into the brain ventricle, MS-222 (Merck) anesthetized 48hpf zebrafish embryos were laterally mounted in 1% low gelling agarose (Merck). Reaction mix containing 0.1U/μL heparinase I and 70,000 MW Dextran-Tetramethylrhodamine (ThermoFisher Scientific) was injected into the 4th ventricle of immobilized embryo. Injected embryos were freed from agarose and allowed to recover in glass bottomed dishes prior to imaging.

## References

Anne, S. L., Govek, E.-E., Ayrault, O., Kim, J. H., Zhu, X., Murphy, D. A., Van Aelst, L., Roussel, M. F., & Hatten, M. E. (2013). WNT3 Inhibits Cerebellar Granule Neuron Progenitor Proliferation and Medulloblastoma Formation via MAPK Activation. PLOS ONE, 8(11), e81769. https://doi.org/10.1371/journal.pone.0081769

Azbazdar, Y., Ozalp, O., Sezgin, E., Veerapathiran, S., Duncan, A. L., Sansom, M. S. P., Eggeling, C., Wohland, T., Karaca, E., & Ozhan, G. (2019). More Favorable Palmitic Acid Over Palmitoleic Acid Modification of Wnt3 Ensures Its Localization and Activity in Plasma Membrane Domains. Frontiers in Cell and Developmental Biology, 7(November), 1–15. https://doi.org/10.3389/fcell.2019.00281

Baeg, G. H., Selva, E. M., Goodman, R. M., Dasgupta, R., & Perrimon, N. (2004). The Wingless morphogen gradient is established by the cooperative action of Frizzled and Heparan Sulfate Proteoglycan receptors. Developmental Biology, 276(1), 89–100. https://doi.org/10.1016/j.ydbio.2004.08.023

Balciunas, D., Wangensteen, K. J., Wilber, A., Bell, J., Geurts, A., Sivasubbu, S., Wang, X., Hackett, P. B., Largaespada, D. A., McIvor, R. S., & Ekker, S. C. (2006). Harnessing a high cargo-capacity transposon for genetic applications in vertebrates. PLoS Genetics, 2(11), 1715–1724. https://doi.org/10.1371/journal.pgen.0020169

Bilić, J., Huang, Y. L., Davidson, G., Zimmermann, T., Cruciat, C. M., Bienz, M., & Niehrs, C. (2007). Wnt induces LRP6 signalosomes and promotes dishevelled-dependent LRP6 phosphorylation. Science, 316(5831), 1619–1622. https://doi.org/10.1126/science.1137065

Bulfone, A., Puelles, L., Porteus, M. H., Frohman, M. A., Martin, G. R., & Rubenstein, J. L. R. (1993). Spatially restricted expression of Dlx-1, Dlx-2 (Tes-1), Gbx-2, and Wnt-3 in the embryonic day 12.5 mouse forebrain defines potential transverse and longitudinal segmental boundaries. Journal of Neuroscience, 13(7), 3155–3172. https://doi.org/10.1523/jneurosci.13-07-03155.1993

Callejo, A., Bilioni, A., Mollica, E., Gorfinkiel, N., Andrés, G., Ibáñez, C., Torroja, C., Doglio, L., Sierra, J., & Guerrero, I. (2011). Dispatched mediates Hedgehog basolateral release to form the long-range morphogenetic gradient in the *Drosophila* wing disk epithelium. Proceedings of the National Academy of Sciences, 108(31), 12591 LP – 12598. https://doi.org/10.1073/pnas.1106881108

Chizhikov, V. V., Lindgren, A. G., Currle, D. S., Rose, M. F., Monuki, E. S., & Millen, K. J. (2006). The roof plate regulates cerebellar cell-type specification and proliferation. Development, 133(15), 2793–2804. https://doi.org/10.1242/dev.02441

Chizhikov, V. V., & Millen, K. J. (2005). Roof plate-dependent patterning of the vertebrate dorsal central nervous system. Developmental Biology, 277(2), 287–295. https://doi.org/10.1016/j.ydbio.2004.10.011

Christianson, H. C., & Belting, M. (2014). Heparan sulfate proteoglycan as a cell-surface endocytosis receptor. Matrix Biology, 35, 51–55. https://doi.org/10.1016/j.matbio.2013.10.004

Chu, M. L. H., Ahn, V. E., Choi, H. J., Daniels, D. L., Nusse, R., & Weis, W. I. (2013). Structural studies of wnts and identification of an LRP6 binding site. Structure, 21(7), 1235–1242. https://doi.org/10.1016/j.str.2013.05.006

Clements, W. K., Ong, K. G., & Traver, D. (2009). Zebrafish wnt3 is expressed in developing neural tissue. Developmental Dynamics, 238(7), 1788–1795. https://doi.org/10.1002/dvdy.21977

Clevers, H., & Nusse, R. (2012). Wnt/β-catenin signaling and disease. Cell, 149(6), 1192–1205. https://doi.org/10.1016/j.cell.2012.05.012

Coudreuse, D., & Korswagen, H. C. (2007). The making of Wnt: New insights into Wnt maturation, sorting and secretion. Development, 134(1), 3–12. https://doi.org/10.1242/dev.02699

Dijksterhuis, J. P., Baljinnyam, B., Stanger, K., Sercan, H. O., Ji, Y., Andres, O., Rubin, J. S., Hannoush, R. N., & Schulte, G. (2015). Systematic mapping of WNT-FZD protein interactions reveals functional selectivity by distinct WNT-FZD pairs. Journal of Biological Chemistry, 290(11), 6789–6798. https://doi.org/10.1074/jbc.M114.612648

Enderlein, J., Gregor, I., Patra, D., Dertinger, T., & Kaupp, U. B. (2005). Performance of Fluorescence Correlation Spectroscopy for Measuring Diffusion and Concentration. ChemPhysChem, 6(11), 2324–2336. https://doi.org/10.1002/cphc.200500414

Esteve, P., Sandonìs, A., Ibañez, C., Shimono, A., Guerrero, I., & Bovolenta, P. (2011). Secreted frizzled-related proteins are required for Wnt/β-catenin signalling activation in the vertebrate optic cup. Development, 138(19), 4179–4184. https://doi.org/10.1242/dev.065839

Eugster, C., Panáková, D., Mahmoud, A., & Eaton, S. (2007). Lipoprotein-Heparan Sulfate Interactions in the Hh Pathway. Developmental Cell, 13(1), 57–71. https://doi.org/10.1016/j.devcel.2007.04.019

Foo, Y. H., Naredi-Rainer, N., Lamb, D. C., Ahmed, S., & Wohland, T. (2012). Factors affecting the quantification of biomolecular interactions by fluorescence cross-correlation spectroscopy. Biophysical Journal, 102(5), 1174–1183. https://doi.org/10.1016/j.bpj.2012.01.040

Fuerer, C., Habib, S. J., & Nusse, R. (2010). A study on the interactions between heparan sulfate proteoglycans and Wnt proteins. Developmental Dynamics, 239(1), 184–190. https://doi.org/10.1002/dvdy.22067

Galli, L. M., Barnes, T. L., Secrest, S. S., Kadowaki, T., & Burrus, L. W. (2007). Porcupine-mediated lipid-modification regulates the activity and distribution of Wnt proteins in the chick neural tube. Development, 134(18), 3339–3348. https://doi.org/10.1242/dev.02881

Garriock, R. J., Warkman, A. S., Meadows, S. M., D’Agostino, S., & Krieg, P. A. (2007). Census of vertebrate Wnt genes: Isolation and developmental expression of Xenopus Wnt2, Wnt3, Wnt9a, Wnt9b, Wnt10a, and Wnt16. Developmental Dynamics, 236(5), 1249–1258. https://doi.org/10.1002/dvdy.21156

Gibbs, H. C., Chang-Gonzalez, A., Hwang, W., Yeh, A. T., & Lekven, A. C. (2017). Midbrain-hindbrain boundary morphogenesis: At the intersection of wnt and Fgf signaling. Frontiers in Neuroanatomy, 11(August), 1–17. https://doi.org/10.3389/fnana.2017.00064

Greco, V., Hannus, M., & Eaton, S. (2001). Argosomes: A potential vehicle for the spread of morphogens through epithelia. Cell, 106(5), 633–645. https://doi.org/10.1016/S0092-8674(01)00484-6

Haas, P., & Gilmour, D. (2006). Chemokine Signaling Mediates Self-Organizing Tissue Migration in the Zebrafish Lateral Line. Developmental Cell, 10(5), 673–680. https://doi.org/10.1016/j.devcel.2006.02.019

Herr, P., & Basler, K. (2012). Porcupine-mediated lipidation is required for Wnt recognition by Wls. Developmental Biology, 361(2), 392–402. https://doi.org/10.1016/j.ydbio.2011.11.003

Hikasa, H., & Sokol, S. Y. (2013). Wnt signaling in vertebrate axis specification. Cold Spring Harbor Perspectives in Biology, 5(1). https://doi.org/10.1101/cshperspect.a007955

Hirai, H., Matoba, K., Mihara, E., Arimori, T., & Takagi, J. (2019). Crystal structure of a mammalian Wnt–frizzled complex. Nature Structural and Molecular Biology, 26(May). https://doi.org/10.1038/s41594-019-0216-z

Holzer, T., Liffers, K., Rahm, K., Trageser, B., Özbek, S., & Gradl, D. (2012). Live imaging of active fluorophore labelled Wnt proteins. FEBS Letters, 586(11), 1638–1644. https://doi.org/10.1016/j.febslet.2012.04.035

Hsieh, J. C., Rattner, A., Smallwood, P. M., & Nathans, J. (1999). Biochemical characterization of Wnt-frizzled interactions using a soluble, biologically active vertebrate Wnt protein. Proceedings of the National Academy of Sciences of the United States of America, 96(7), 3546–3551. https://doi.org/10.1073/pnas.96.7.3546

Huang, H., & Kornberg, T. B. (2015). Myoblast cytonemes mediate Wg signaling from the wing imaginal disc and Delta-Notch signaling to the air sac primordium. ELife, 4(MAY), 1–22. https://doi.org/10.7554/eLife.06114

Janda, C. Y., Waghray, D., Levin, A. M., Thomas, C., & Garcia, K. C. (2012). Structural Basis of Wnt. Science, 337(July), 59–64. https://doi.org/10.1126/science.1222879.Structural

Jessell TM. (2000). Neuronal specification in the spinal cord:inductive signals and transcriptional codes. Nature Reviews Genetics, 1(October), 20–29.

Kaiser, K., Gyllborg, D., Procházka, J., Salašová, A., Kompaníková, P., Molina, F. L., Laguna-Goya, R., Radaszkiewicz, T., Harnoš, J., Procházková, M., Potěšil, D., Barker, R. A., Casado, Á. G., Zdráhal, Z., Sedláček, R., Arenas, E., Villaescusa, J. C., & Bryja, V. (2019). WNT5A is transported via lipoprotein particles in the cerebrospinal fluid to regulate hindbrain morphogenesis. Nature Communications, 10(1), 1–15. https://doi.org/10.1038/s41467-019-09298-4

Kang, M., Cordova, M., Chen, C. S. J., & Rajadhyaksha, M. (2015). Simplified equation to extract diffusion coefficients from confocal FRAP data. Journal of Investigative Dermatology, 135(2), 612–615. https://doi.org/10.1038/jid.2014.371

Kapusta, P., Macháň, R., Benda, A., & Hof, M. (2012). Fluorescence Lifetime Correlation Spectroscopy (FLCS): concepts, applications and outlook. International Journal of Molecular Sciences, 13(10), 12890–12910. https://doi.org/10.3390/ijms131012890

Kim, S. A., Heinze, K. G., & Schwille, P. (2007). Fluorescence correlation spectroscopy in living cells. Nature Methods, 4(11), 963–973. https://doi.org/10.1038/nmeth1104

Kimmel, C. B., Ballard, W. W., Kimmel, S. R., Ullmann, B., & Schilling, T. F. (1995). Stages of embryonic development of the zebrafish. Developmental Dynamics : An Official Publication of the American Association of Anatomists, 203(3), 253–310. https://doi.org/10.1002/aja.1002030302

Kirkpatrick, C. A., & Selleck, S. B. (2007). Heparan sulfate proteoglycans at a glance. Journal of Cell Science, 120(11), 1829–1832. https://doi.org/10.1242/jcs.03432

Klonis, N., Rug, M., Harper, I., Wickham, M., Cowman, A., & Tilley, L. (2002). Fluorescence photobleaching analysis for the study of cellular dynamics. European Biophysics Journal, 31(1), 36–51. https://doi.org/10.1007/s00249-001-0202-2

Kondrychyn, I., Teh, C., Sin, M., & Korzh, V.. -001216069090203U-main. pd. (2013). Stretching Morphogenesis of the Roof Plate and Formation of the Central Canal. PLoS ONE, 8(2), 1–12. https://doi.org/10.1371/journal.pone.0056219

Koppel, D. E., Axelrod, D., Schlessinger, J., Elson, E. L., & Webb, W. W. (1976). Dynamics of fluorescence marker concentration as a probe of mobility. Biophysical Journal, 16(11), 1315–1329. https://doi.org/10.1016/S0006-3495(76)85776-1

Krichevsky, O., & Bonnet, G. (2002). Fluorescence correlation spectroscopy: the technique and its applications. Reports on Progress in Physics, 65(2), 251–297. https://doi.org/10.1088/0034-4885/65/2/203

Li, Q., Lau, A., Morris, T. J., Guo, L., Fordyce, C. B., & Stanley, E. F. (2004). A Syntaxin 1, Gαo, and N-Type Calcium Channel Complex at a Presynaptic Nerve Terminal: Analysis by Quantitative Immunocolocalization. Journal of Neuroscience, 24(16), 4070–4081. https://doi.org/10.1523/JNEUROSCI.0346-04.2004

Lipinski, C., & Hopkins, A. (2004). Navigating chemical space for biology and medicine. Nature, 432(7019), 855–861. https://doi.org/10.1038/nature03193

Liu, P., Wakamiya, M., Shea, M. J., Albrecht, U., Behringer, R. R., & Bradley, A. (1999). Requirement for Wnt3 in vertebrate axis formation. Nature Genetics, 22(4), 361–365. https://doi.org/10.1038/11932

Logan, C. Y., & Nusse, R. (2004). the Wnt Signaling Pathway in Development and Disease. Annual Review of Cell and Developmental Biology, 20(1), 781–810. https://doi.org/10.1146/annurev.cellbio.20.010403.113126

Lord, N. D., Carte, A. N., Abitua, P. B., & Schier, A. F. (2019). The pattern of Nodal morphogen signaling is shaped by co-receptor expression. BioRxiv, 2019.12.30.891101. https://doi.org/10.1101/2019.12.30.891101

Macháň, R., Foo, Y. H., & Wohland, T. (2016). On the Equivalence of FCS and FRAP: Simultaneous Lipid Membrane Measurements. Biophysical Journal, 111(1), 152–161. https://doi.org/10.1016/j.bpj.2016.06.001

Magde, D., Elson, E. L., & Webb, W. W. (1974). Fluorescence correlation spectroscopy. II. An experimental realization. Biopolymers, 13(1), 29–61. https://doi.org/10.1002/bip.1974.360130103

Mattes, B., Dang, Y., Greicius, G., Kaufmann, L. T., Prunsche, B., Rosenbauer, J., Stegmaier, J., Mikut, R., Özbek, S., Nienhaus, G. U., Schug, A., Virshup, D. M., & Scholpp, S. (2018). Wnt/PCP controls spreading of Wnt/β-catenin signals by cytonemes in vertebrates. ELife, 7, 1–28. https://doi.org/10.7554/eLife.36953

Mihara, E., Hirai, H., Yamamoto, H., Tamura-Kawakami, K., Matano, M., Kikuchi, A., Sato, T., & Takagi, J. (2016). Active and water-soluble form of lipidated wnt protein is maintained by a serum glycoprotein afamin/α-albumin. ELife, 5(FEBRUARY2016), 1–19. https://doi.org/10.7554/eLife.11621

Mii, Y., & Taira, M. (2009). Secreted Frizzled-related proteins enhance the diffusion of Wnt ligands and expand their signalling range. Development, 136(24), 4083–4088. https://doi.org/10.1242/dev.032524

Mii, Y., Yamamoto, T., Takada, R., Mizumoto, S., Matsuyama, M., Yamada, S., Takada, S., & Taira, M. (2017). Roles of two types of heparan sulfate clusters in Wnt distribution and signaling in Xenopus. Nature Communications, 8(1), 1–19. https://doi.org/10.1038/s41467-017-02076-0

Mikels, A. J., & Nusse, R. (2006). Wnts as ligands: Processing, secretion and reception. Oncogene, 25(57), 7461–7468. https://doi.org/10.1038/sj.onc.1210053

Moon, R. T., Bowerman, B., Boutros, M., & Perrimon, N. (2002). The promise and perils of Wnt signaling through β-catenin. Science, 296(5573), 1644–1646. https://doi.org/10.1126/science.1071549

Müller, P., Rogers, K. W., Jordan, B. M., Lee, J. S., Robson, D., Ramanathan, S., & Schier, A. F. (2012). Differential diffusivity of nodal and lefty underlies a reaction-diffusion patterning system. Science, 336(6082), 721–724. https://doi.org/10.1126/science.1221920

Müller, P., Rogers, K. W., Yu, S. R., Brand, M., & Schier, A. F. (2013). Morphogen transport. Development, 140(8), 1621–1638. https://doi.org/10.1242/dev.083519

Mulligan, K. A., Fuerer, C., Ching, W., Fish, M., Willert, K., & Nusse, R. (2012). Secreted Wingless-interacting molecule (Swim) promotes long-range signaling by maintaining Wingless solubility. Proceedings of the National Academy of Sciences of the United States of America, 109(2), 370–377. https://doi.org/10.1073/pnas.1119197109

Naschberger, A., Orry, A., Lechner, S., Bowler, M. W., Nurizzo, D., Novokmet, M., Keller, M. A., Oemer, G., Seppi, D., Haslbeck, M., Pansi, K., Dieplinger, H., & Rupp, B. (2017). Structural Evidence for a Role of the Multi-functional Human Glycoprotein Afamin in Wnt Transport. Structure, 25(12), 1907-1915.e5. https://doi.org/10.1016/j.str.2017.10.006

Neumann, S., Coudreuse, D. Y. M., Van Der Westhuyzen, D. R., Eckhardt, E. R. M., Korswagen, H. C., Schmitz, G., & Sprong, H. (2009). Mammalian Wnt3a is released on lipoprotein particles. Traffic, 10(3), 334–343. https://doi.org/10.1111/j.1600-0854.2008.00872.x

Ng, X. W., Teh, C., Korzh, V., & Wohland, T. (2016). The Secreted Signaling Protein Wnt3 Is Associated with Membrane Domains In Vivo: A SPIM-FCS Study. Biophysical Journal, 111(2), 418–429. https://doi.org/10.1016/j.bpj.2016.06.021

Niehrs, C. (2012). The complex world of WNT receptor signalling. Nature Reviews Molecular Cell Biology, 13(12), 767–779. https://doi.org/10.1038/nrm3470

Ohkawara, B., Yamamoto, T. S., Tada, M., & Ueno, N. (2003). Role of glypican 4 in the regulation of convergent extension movements during gastrulation in Xenopus laevis. Development, 130(10), 2129–2138. https://doi.org/10.1242/dev.00435

Padilla-Parra, S., Audugé, N., Coppey-Moisan, M., & Tramier, M. (2011). Dual-color fluorescence lifetime correlation spectroscopy to quantify protein–protein interactions in live cell. Microscopy Research and Technique, 74(8), 788–793. https://doi.org/10.1002/jemt.21015

Panáková, D., Sprong, H., Marois, E., Thiele, C., & Eaton, S. (2005). Lipoprotein particles are required for Hedgehog and Wingless signalling. Nature, 435(7038), 58–65. https://doi.org/10.1038/nature03504

Raible, F., & Brand, M. (2004). Divide et Impera–the midbrain–hindbrain boundary and its organizer. Trends in Neurosciences, 27(12), 727–734.

Ries, J., Yu, S. R., Burkhardt, M., Brand, M., & Schwille, P. (2009). Modular scanning FCS quantifies receptor-ligand interactions in living multicellular organisms. Nature Methods, 6(9), 643–645. https://doi.org/10.1038/nmeth.1355

Routledge, D., & Scholpp, S. (2019). Mechanisms of intercellular Wnt transport. Development, 146(10), dev176073. https://doi.org/10.1242/dev.176073

Saied-Santiago, K., Townley, R. A., Attonito, J. D., Cunha, D. S. da, Díaz-Balzac, C. A., Tecl, E., & Bülow, H. E. (2017). Coordination of Heparan Sulfate Proteoglycans with Wnt Signaling To Control Cellular Migrations and. 206(August), 1951–1967. https://doi.org/10.1534/genetics.116.198739/-/DC1.1

Schubert, M., & Holland, L. Z. (2013). The Wnt gene family and the evolutionary conservation of Wnt expression. In Madame Curie Bioscience Database. Landes Bioscience.

Schulte, G., & Wright, S. C. (2018). Frizzleds as GPCRs – More Conventional Than We Thought! Trends in Pharmacological Sciences, 0(0), 1–15. https://doi.org/10.1016/j.tips.2018.07.001

Schwille, P., Meyer-Almes, F. J., & Rigler, R. (1997). Dual-color fluorescence cross-correlation spectroscopy for multicomponent diffusional analysis in solution. Biophysical Journal, 72(4), 1878–1886. https://doi.org/10.1016/S0006-3495(97)78833-7

Sezgin, E., Azbazdar, Y., Ng, X. W., Teh, C., Simons, K., Weidinger, G., Wohland, T., Eggeling, C., & Ozhan, G. (2017). Binding of canonical Wnt ligands to their receptor complexes occurs in ordered plasma membrane environments. FEBS Journal, 284(15), 2513–2526. https://doi.org/10.1111/febs.14139

Shi, X., Yong, H. F., Sudhaharan, T., Chong, S. W., Korzh, V., Ahmed, S., & Wohland, T. (2009). Determination of dissociation constants in living zebrafish embryos with single wavelength fluorescence cross-correlation spectroscopy. Biophysical Journal, 97(2), 678–686. https://doi.org/10.1016/j.bpj.2009.05.006

Speer, K. F., Sommer, A., Tajer, B., Mullins, M. C., Klein, P. S., & Lemmon, M. A. (2019). Non-acylated Wnts Can Promote Signaling. Cell Reports, 26(4), 875-883.e5. https://doi.org/10.1016/j.celrep.2018.12.104

Stanganello, E., Hagemann, A. I. H., Mattes, B., Sinner, C., Meyen, D., Weber, S., Schug, A., Raz, E., & Scholpp, S. (2015). Filopodia-based Wnt transport during vertebrate tissue patterning. Nature Communications, 6, 1–14. https://doi.org/10.1038/ncomms6846

Sun, G., Guo, S.-M., Teh, C., Korzh, V., Bathe, M., & Wohland, T. (2015). Bayesian Model Selection Applied to the Analysis of Fluorescence Correlation Spectroscopy Data of Fluorescent Proteins *in Vitro* and *in Vivo*. Analytical Chemistry, 87(8), 4326–4333. https://doi.org/10.1021/acs.analchem.5b00022

Tao, Q., Yokota, C., Puck, H., Kofron, M., Birsoy, B., Yan, D., Asashima, M., Wylie, C. C., Lin, X., & Heasman, J. (2005). Maternal Wnt11 activates the canonical Wnt signaling pathway required for axis formation in Xenopus embryos. Cell, 120(6), 857–871. https://doi.org/10.1016/j.cell.2005.01.013

Teh, C., Sun, G., Shen, H., Korzh, V., & Wohland, T. (2015). Modulating the expression level of secreted Wnt3 influences cerebellum development in zebrafish transgenics. Development, 142(21), 3721–3733. https://doi.org/10.1242/dev.127589

Topczewski, J., Sepich, D. S., Myers, D. C., Walker, C., Amores, A., Lele, Z., Hammerschmidt, M., Postlethwait, J., & Solnica-Krezel, L. (2001). The Zebrafish Glypican Knypek Controls Cell Polarity during Gastrulation Movements of Convergent Extension. Developmental Cell, 1(2), 251–264. https://doi.org/10.1016/S1534-5807(01)00005-3

Veerapathiran, S., & Wohland, T. (2018). Fluorescence techniques in developmental biology. Journal of Biosciences, 43(3), 541–553. https://doi.org/10.1007/s12038-018-9768-z

Wang, Y., Wang, X., Wohland, T., & Sampath, K. (2016). Extracellular interactions and ligand degradation shape the nodal morphogen gradient. ELife, 5(APRIL2016), 1–19. https://doi.org/10.7554/eLife.13879

Willems, B., Tao, S., Yu, T., Huysseune, A., Witten, P. E., & Winkler, C. (2015). The Wnt Co-Receptor Lrp5 Is Required for Cranial Neural Crest Cell Migration in Zebrafish. PLoS ONE, 1–21. https://doi.org/10.1371/journal.pone.0131768

Wilsie, L. C., & Orlando, R. A. (2003). The low density lipoprotein receptor-related protein complexes with cell surface heparan sulfate proteoglycans to regulate proteoglycan-mediated lipoprotein catabolism. Journal of Biological Chemistry, 278(18), 15758–15764. https://doi.org/10.1074/jbc.M208786200

Wilson, M. A. (2017). Structural Insight into a Fatty-Acyl Chaperone for Wnt Proteins. Structure, 25(12), 1781–1782. https://doi.org/10.1016/j.str.2017.11.009

Wu, C. H., & Nusse, R. (2002). Ligand receptor interactions in the Wnt signaling pathway in Drosophila. Journal of Biological Chemistry, 277(44), 41762–41769. https://doi.org/10.1074/jbc.M207850200

Wurst, W., & Bally-Cuif, L. (2001). Neural plate patterning: upstream and downstream of the isthmic organizer. Nature Reviews Neuroscience, 2(2), 99–108.

Yavas, S., Macháň, R., & Wohland, T. (2016). The Epidermal Growth Factor Receptor Forms Location-Dependent Complexes in Resting Cells. Biophysical Journal, 111(10), 2241–2254. https://doi.org/10.1016/j.bpj.2016.09.049

Zhu, S., Welsch, R. E., & Matsudaira, P. T. (2016). A method to quantify co-localization in biological images. Proceedings of the Annual International Conference of the IEEE Engineering in Medicine and Biology Society, EMBS, 2016-Octob, 3887–3890. https://doi.org/10.1109/EMBC.2016.7591577

