## Supplementary Figures for "Wnt3 distribution in the zebrafish brain is determined by expression, diffusion and multiple molecular interactions"

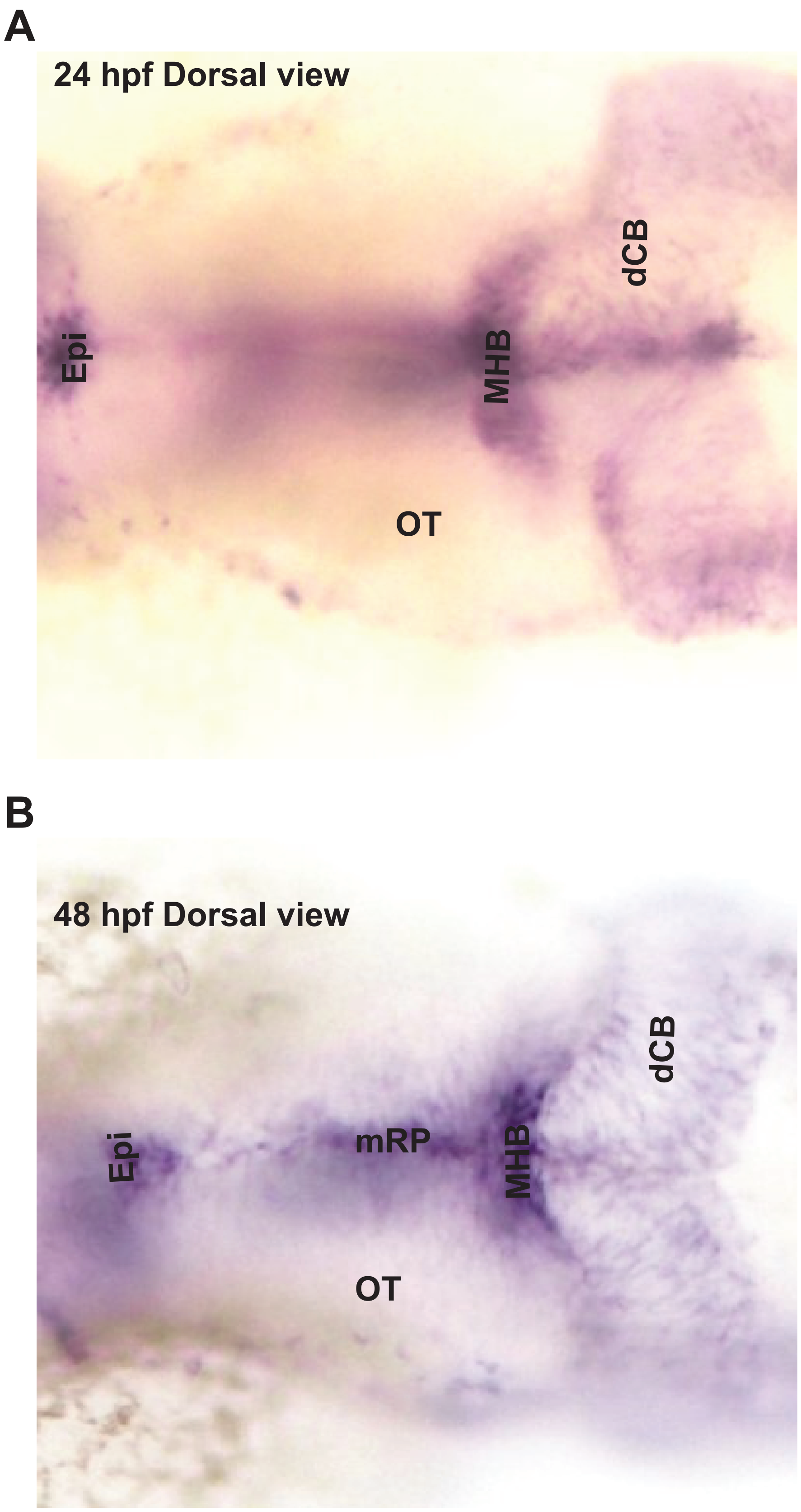


**Supplementary Figure 1: Expression of *wnt3* transcript by whole mount in situ hybridization.** In situ hybridization detects expression of wnt3 at **(A)** 24 hpf and **(B)** 48 hpf. Ce, cerebellum; Epi, Epithalamus; MHB, midbrain-hindbrain boundary; mRP, midbrain roof plate; OT, optic tectum.

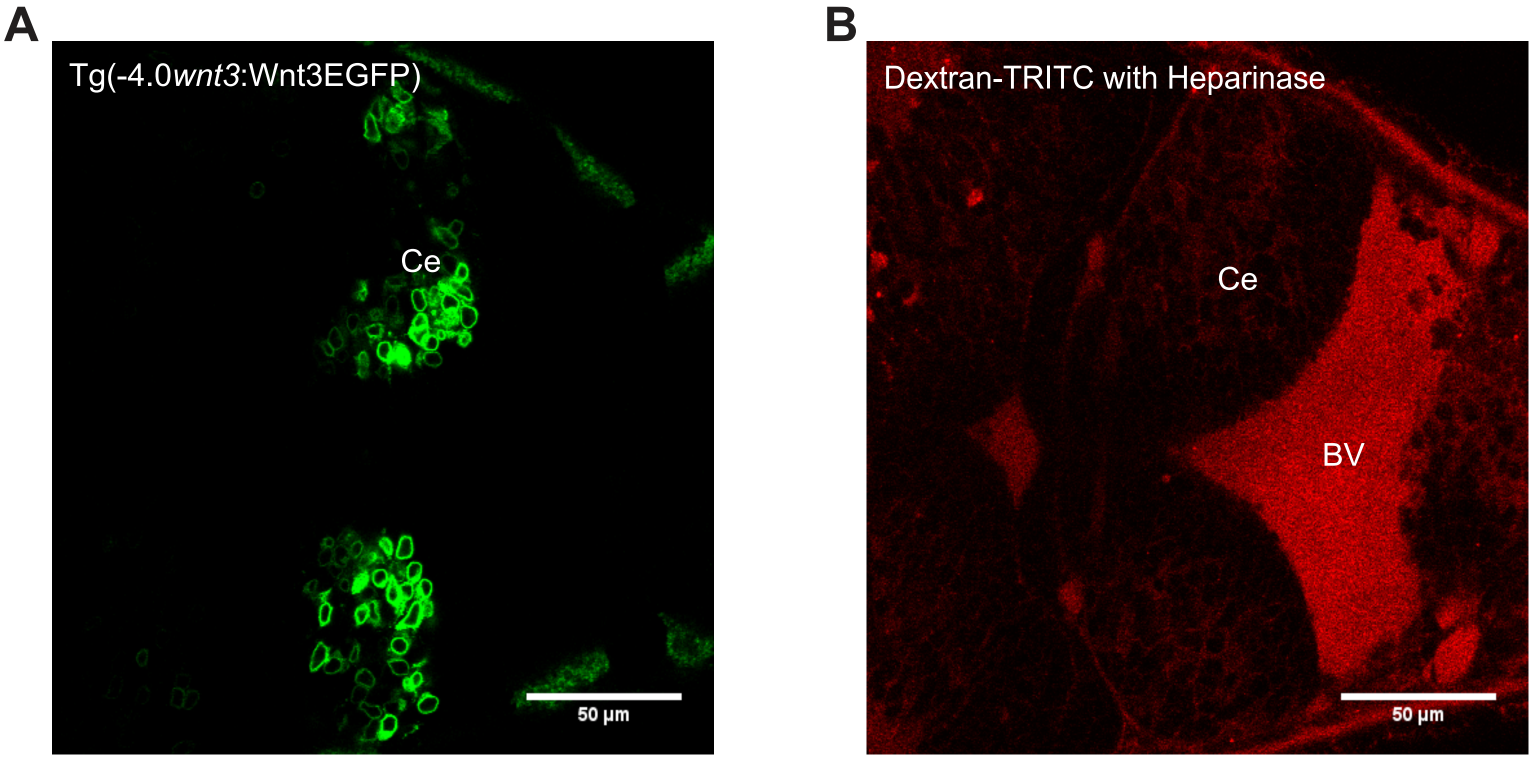


**Supplementary Figure 2: Tg(-4.0*wnt3*:Wnt3EGFP) embryos treated by Heparinase.**
**(A)** The expression of Wnt3EGFP after heparinase treatment. **(B)** Distribution of Dextran-TRITC coinjected with Heparinase in the brain ventricle of Tg(-4.0*wnt3*:Wnt3EGFP) embryo. BV, Brain ventricle; Ce, cerebellum. Scale bar 50 µm.

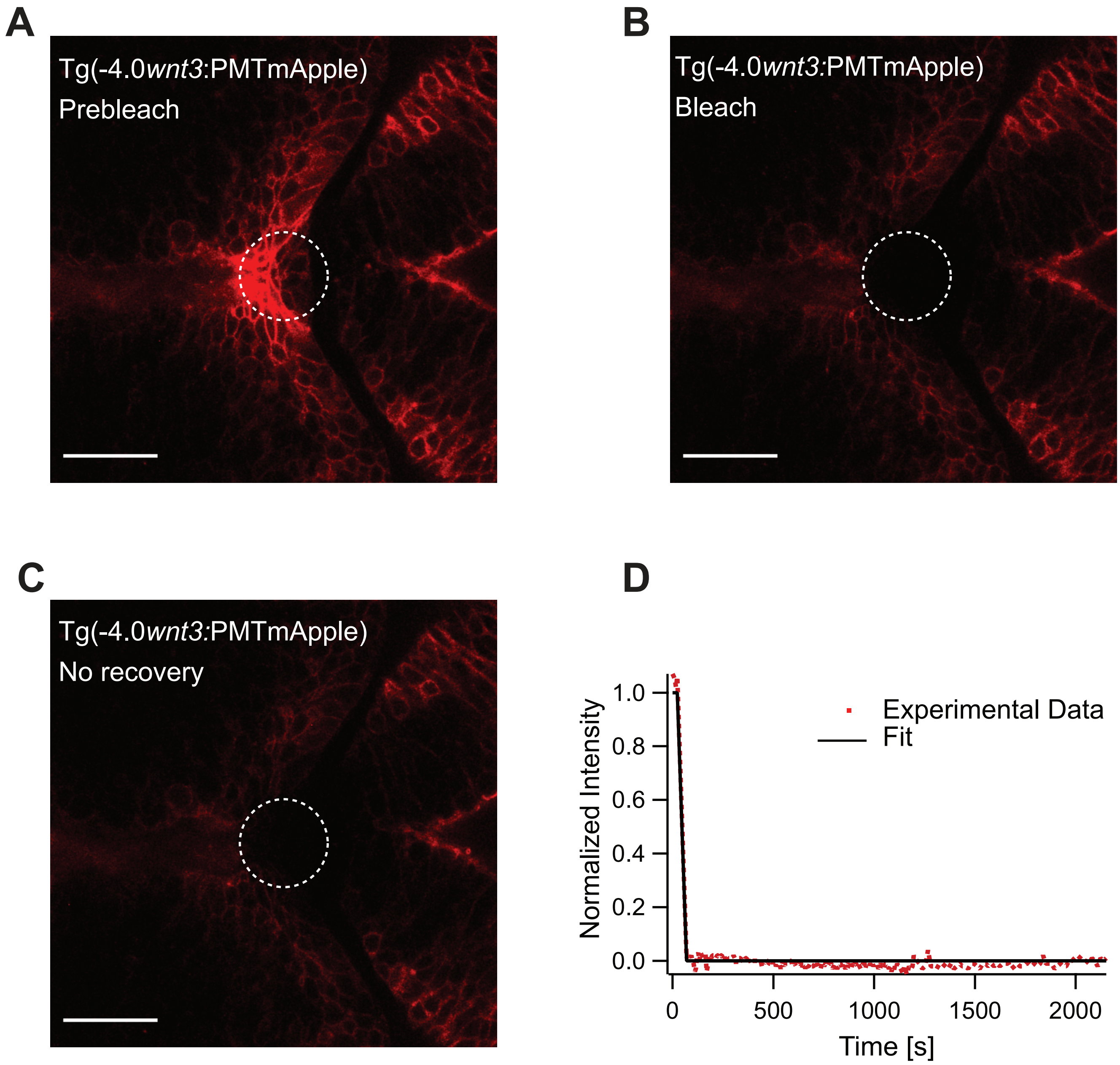


**Supplementary Figure 3: Representative fluorescence recovery of PMTmApple after photobleaching.** **(A)** Expression of PMTmApple in Tg(-4.0*wnt3*:PMTmApple) before photobleaching. **(B)** Photobleached region of PMTmApple. **(C)** No recovery of fluorescence intensity in the bleached region as PMT re-mains tethered to the cell membrane and does not diffuse. **(D)** Fluorescence recovery curve for PMTmApple. Scale bar 30 µm.

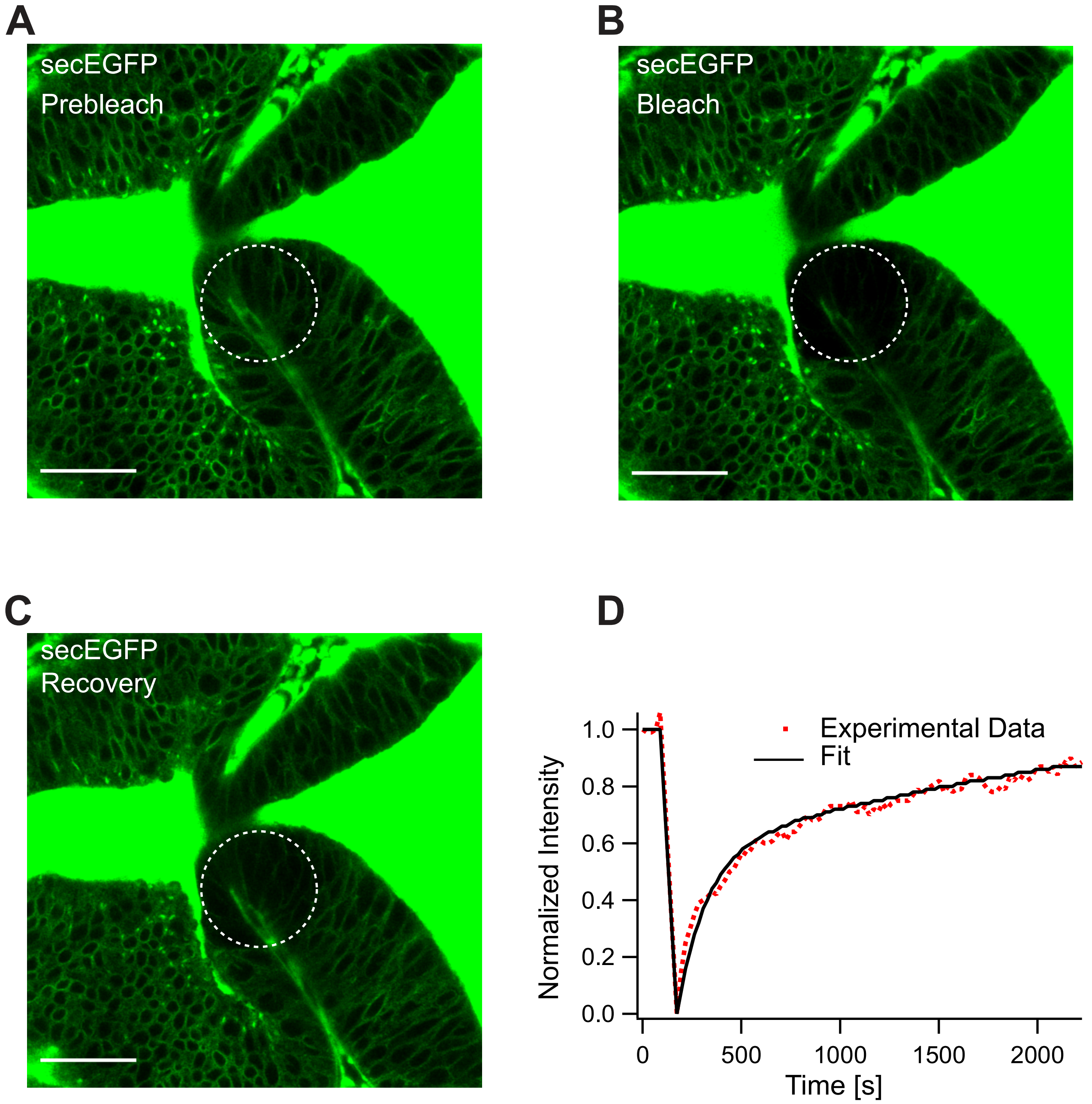


**Supplementary Figure 4: Representative fluorescence recovery of secEGFP after photobleaching.** **(A)** Expression of secEGFP before photobleaching. **(B)** Photobleached region of secEGFP. **(C)** Recovery of fluorescence intensity in the bleached region due to diffusion of molecules from the neighboring un-bleached regions. **(D)** Fluorescence recovery curve for secEGFP with a quick recovery time ($\tau_{fast}$) of ~ 45 s and a fraction of mobile component (Fm) of ~ 0.85. The average global diffusion coefficient ($D_{eff}$) measured for secEGFP was 13 ± 4 μm$^2$/s (N=10). Scale bar 30 μm.

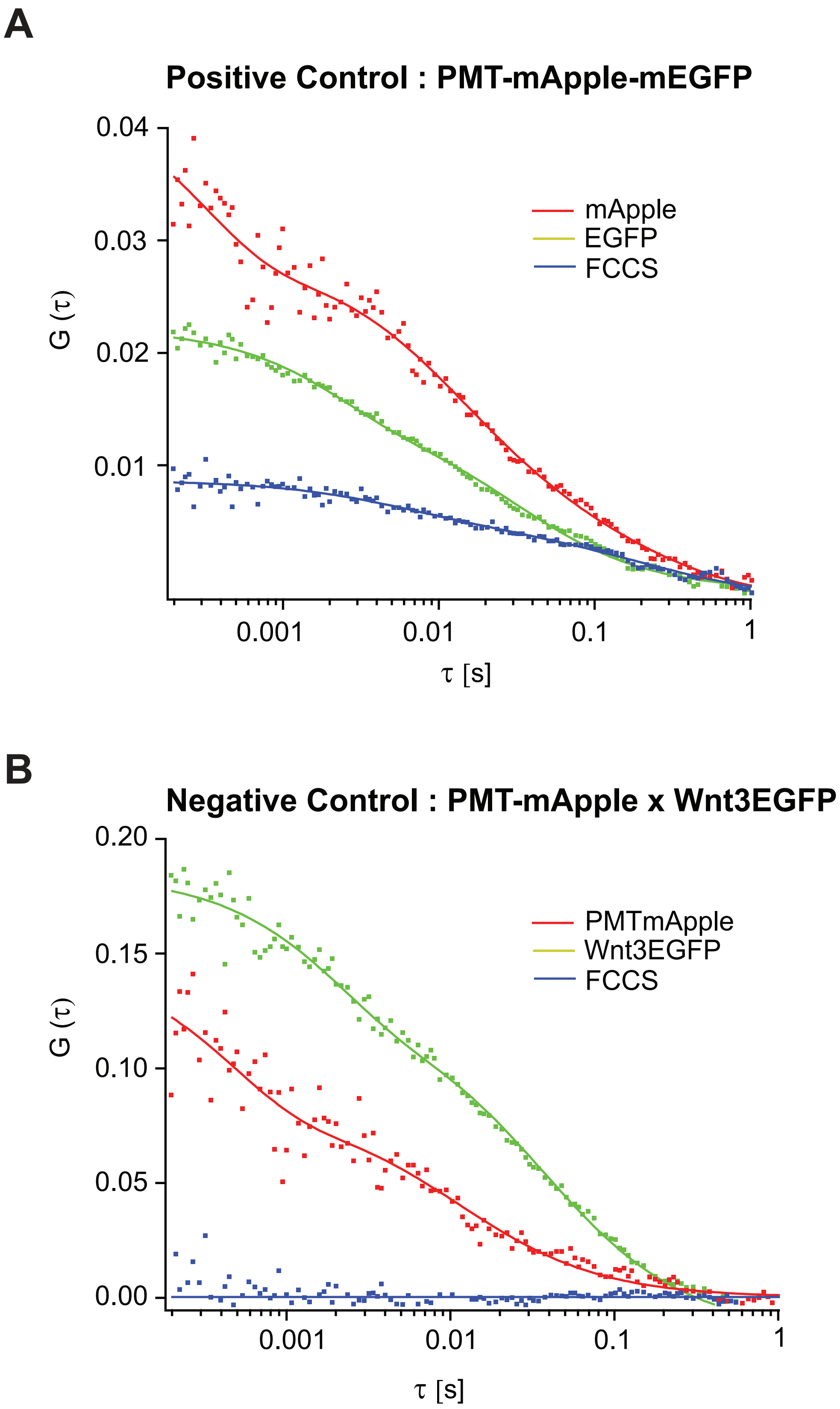


**Supplementary Figure 5: Representative FCCS measurements for positive and negative control.** Representative auto- and cross-correlation functions for **(A)** PMT-mApple-mEGFP which is a positive control showing a clear cross-correlation and **(B)** Negative control PMT-mApple and Wnt3-EGFP showing no cross-correlations.

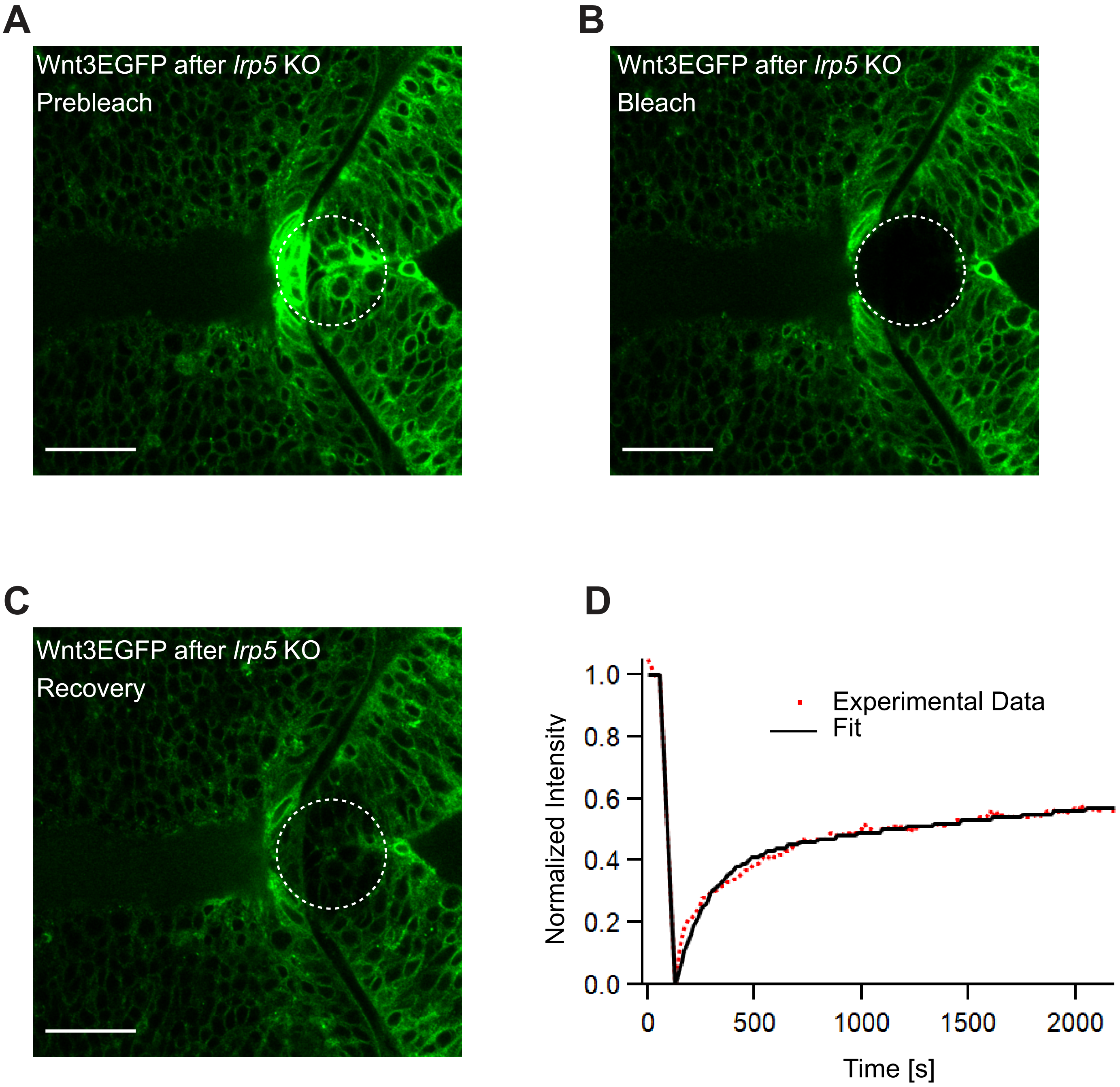


**Supplementary Figure 6: Representative fluorescence recovery of Wnt3EGFP in *lrp5* knocked-down embryos after photobleaching. (A)** Expression of Wnt3EGFP before photobleaching. **(B)** Photobleached region of Wnt3EGFP. **(C)** Recovery of fluorescence intensity in the bleached region due to diffusion of molecules from the neighboring unbleached regions. **(D)** Fluorescence recovery curve for Wnt3EGFP with a recovery time ($\tau_{fast}$) of ~ 150 s and a fraction of mobile component (Fm) of ~ 0.5. The average global diffusion coefficient ($D_{eff}$) measured for Wnt3EGFP after *lrp5* knock-down was 3 ± 0.8 $\mu m^2/s$ (N=10). Scale bar 30 µm.
